# C53 interacting with UFM1-protein ligase 1 regulates microtubule nucleation in response to ER stress

**DOI:** 10.1101/2020.12.23.424116

**Authors:** Anastasiya Klebanovych, Stanislav Vinopal, Eduarda Dráberová, Vladimíra Sládková, Tetyana Sulimenko, Vadym Sulimenko, Věra Vosecká, Libor Macůrek, Agustin Legido, Pavel Dráber

## Abstract

ER distribution depends on microtubules, and ER homeostasis disturbance activates the unfolded protein response resulting in ER remodeling. CDK5RAP3 (C53) implicated in various signaling pathways interacts with UFM1-protein ligase 1 (UFL1), which mediates the ufmylation of proteins in response to ER stress. Here we find that UFL1 and C53 associate with γ-tubulin ring complex proteins. Knockout of *UFL1* or *C53* in human osteosarcoma cells induces ER stress, centrosomal microtubule nucleation accompanied by γ-tubulin accumulation and ER expansion. C53, whose protein level is modulated by UFL1, associates with the centrosome and rescues microtubule nucleation in cells lacking UFL1. Pharmacological induction of ER stress by tunicamycin also leads to increased microtubule nucleation and ER expansion. Furthermore, tunicamycin suppresses the association of C53 with the centrosome. These findings point to a novel mechanism for the relief of ER stress by stimulation of centrosomal microtubule nucleation.

## Introduction

The endoplasmic reticulum consists of a continuous network of membranous sheets and tubules spanning the cytoplasm. It plays critical roles in a wide range of processes, including synthesis, folding, modification, and transport of proteins, synthesis and distribution of lipids, and Ca^2+^ storage. A diverse array of cellular stresses can lead to an imbalance between the protein-folding capacity and protein-folding load (Chakrabarti et al., 2011). The perturbation of ER homeostasis (ER stress) is ameliorated by triggering signaling cascades of the unfolded protein response (UPR), which engage effector mechanisms leading to homeostasis restoration. These mechanisms include increasing the capacity of the ER, increasing the degradation of ER luminal proteins or upregulation of chaperones and luminal protein modifications, or folding enzymes (Smith and Wilkinson, 2017). In the course of ER stress, the ER undergoes rapid and extensive remodeling hallmarked by the expansion of its lumen and an increase in tubules (Schuck et al., 2009). In mammalian cells, ER distribution is dependent on microtubules (Waterman-Storer and Salmon, 1998).

Microtubules, composed of α-and β-tubulin heterodimers, are highly dynamic and display dynamic instability characterized by altering phases of growth and shrinkage. During interphase, microtubules are mainly nucleated at the centrosome (microtubule organizing centers; MTOC) and radiate toward the cell periphery.

γ-Tubulin, a conserved member of the tubulin superfamily, is essential for microtubule nucleation in all eukaryotes (Oakley and Oakley, 1989). Together with other proteins named Gamma-tubulin Complex Proteins (GCPs; GCP2-6), it assembles into γ-Tubulin Ring Complex (γTuRC), which in mammalian cells effectively catalyzes microtubule nucleation. GCP2-6 each bind directly to γ-tubulin and assemble into cone-shaped structure of γTuRC (Kollman et al., 2011; Oakley et al., 2015). Recent high-resolution structural studies revealed details of asymmetric structure of γTuRC (Consolati et al., 2020; Liu et al., 2020; Wieczorek et al., 2020).

The γTuRCs are typically concentrated at MTOCs such as centrosomes and basal bodies in animals. They also associate with cellular membranes, including Golgi apparatus (Chabin-Brion et al., 2001), where they participate in non-centrosomal microtubule nucleation (Oakley et al., 2015). The majority of γTuRCs are generally inactive in the cytosol and become active at MTOCs. The mechanisms of γTuRC activation in cells are not fully understood. Current data suggest that γTuRC can be activated by structural rearrangement of γTuRC, phosphorylation or allosterically by binding to γTuRC tethering or modulating proteins accumulated in MTOCs (Liu et al., 2020; Sulimenko et al., 2017).

Ubiquitin-fold modifier (UFM1) is an ubiquitin-like posttranslational modifier that targets proteins through a process called ufmylation (Komatsu et al., 2004; Tatsumi et al., 2010). Conjugation of UFM1 to proteins is mediated by a process analogous to ubiquitination and requires specific activating (E1), conjugating (E2), and ligating (E3) enzymes. Unlike ubiquitination, the modification of proteins by ubiquitin-like modifiers generally serves as a non-proteolysis signal. It regulates various cellular processes by altering the substrate structure, stability, localization, or protein-protein interactions. The ufmylation has been reported to regulate multiple cellular processes, including the ER stress response, ribosome function, control of gene expression, DNA damage response, and cell differentiation (Gerakis et al., 2019). Whether ufmylation controls microtubule organization is unknown. E3 UFM1-protein ligase 1 (also known as KIAA0776, RCAD, NLBP or Maxer; hereafter denoted as UFL1) (Kwon et al., 2010; Shiwaku et al., 2010; Tatsumi et al., 2010; Wu et al., 2010) is mainly located at the cytosolic side of the ER membrane (Shiwaku et al., 2010), where it is found in complex with DDRGK domain-containing protein 1 (DDRGK1, UFBP1) (Lemaire et al., 2011), the first identified substrate of ufmylation (Tatsumi et al., 2010). UPR directly controls the ufmylation pathway under ER stress at the transcriptional level (Zhang et al., 2012; Zhu et al., 2019).

Putative tumor suppressor CDK5 regulatory subunit-associated protein 3 (also known as CDK5RAP3, LZAP; hereafter denoted as C53) has initially been identified as a binding protein of CDK5 activator (Ching et al., 2000). Subsequent research showed that C53 exerts multiple roles in regulating the cell cycle, DNA damage response, cell survival, cell adherence/invasion, tumorigenesis, and metastasis (Jiang et al., 2005; Jiang et al., 2009; Liu et al., 2011; Wang et al., 2007; Zhao et al., 2011). Moreover, C53 is also involved in UPR (Zhang et al., 2012). Several studies have reported the interactions between C53 and UFL1 (Kwon et al., 2010; Shiwaku et al., 2010; Wu et al., 2010), and it was suggested that C53 could serve as UFL1 substrate adaptor (Yang et al., 2019). On the other hand, UFL1 regulates the stability of both C53 and DDRGK1 (Wu et al., 2010). Reports indicating the involvement of C53 in the modulation of microtubule organization are rare. It has been shown that a peptide from caspase-dependent cleavage of C53 participates in microtubule bundling and rupture of the nuclear envelope in apoptotic cells (Wu et al., 2013). In addition, we have demonstrated that C53 forms complexes with nuclear γ-tubulin and that γ-tubulin antagonizes the inhibitory effect of C53 on G2/M checkpoint activation by DNA damage (Hořejší et al., 2012).

In the present study, we provide evidence that C53, which associates with UFL1 and γTuRC proteins, has an important role in the modulation of centrosomal microtubule nucleation in cells under ER stress. The interaction of ER membranes with newly formed microtubules could promote ER expansion to the cell periphery and help restore ER homeostasis.

## Results

### Identification of UFL1 as γ-tubulin interactor

We have previously reported the intrinsic association of γ-tubulin with cellular membranes in mouse P19 embryonal carcinoma cells undergoing neuronal differentiation (Macurek et al., 2008). To identify the potential interacting partners for membrane-bound γ-tubulin, we performed immunoprecipitation experiments with anti-peptide mAb to γ-tubulin and extracts from the microsomal fraction of differentiated P19 cells. The bound proteins were eluted with the peptide used for immunization or with a negative control peptide. In repeated experiments, a 90-kDa protein was specifically eluted with the immunization peptide but not with the control peptide. The protein was subjected to MALDI/MS fingerprint analysis and identified as UFL1 (UniProtKB identifier O94874) (Table S1). UFL1 mainly associates with the endoplasmic reticulum (ER) membranes (Shiwaku et al., 2010; Tatsumi et al., 2010) and interacts with C53 (Kwon et al., 2010; Wu et al., 2010). We showed that C53 is present in γ-tubulin immunocomplexes from the nuclear fraction of human HeLa S3 cells (Hořejší et al., 2012). Collectively these data indicate that UFL1 and C53 can form complexes containing γ-tubulin.

### UFL1 and C53 associate with γTuRC proteins

To ascertain whether membranous UFL1 associates both with C53 and γTuRC proteins in different cell types, we first performed immunoprecipitation experiments with extracts from the crude membranous fraction (P2) from human osteosarcoma U2OS cells. The fraction comprised UFL1, C53, γ-tubulin, GCP2, and calnexin (a marker of ER) but was devoid of α-tubulin, GM130, and histone H1.4 representing, cytosolic, Golgi apparatus and nuclear proteins, respectively (Fig. S1A, lane 5). Using two anti-UFL1 Abs recognizing epitopes in distinct UFL1 aa sequence regions (301-389 [UFL1_301-389_] and 438-793 [UFL1_438-793_]) and Abs to C53, γ-tubulin, and GCP2 for reciprocal precipitations, we revealed association of UFL1 and C53 with γTuRC proteins (Fig. 1A; Fig. S1B). As expected, we also verified the association of C53 with UFL1 (Fig. 1A, b and e).

**Figure 1.**
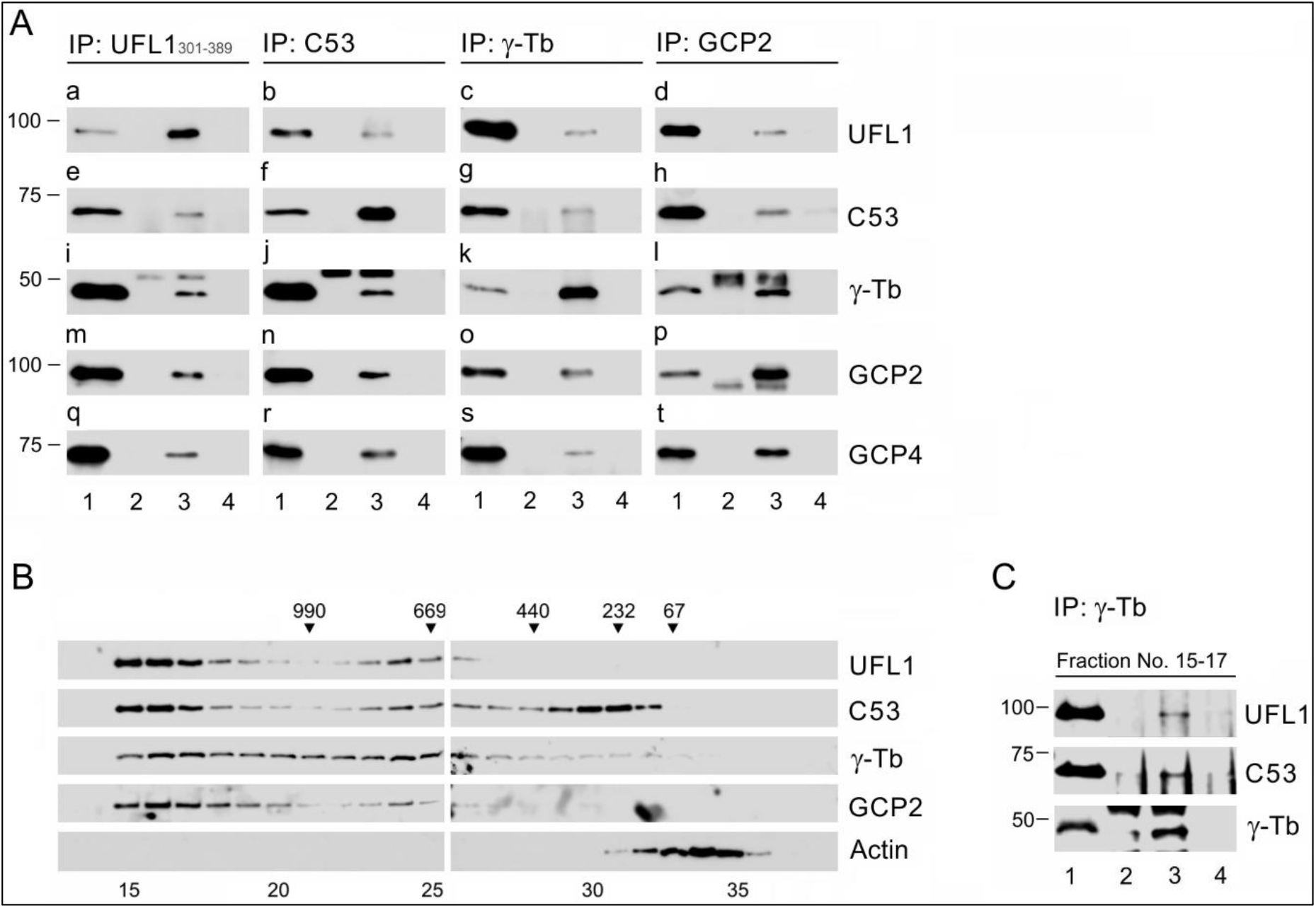
UFL1 and C53 interact with γTuRC proteins. (**A**) Immunoprecipitation experiments. Extracts from the membranous fraction (P2) of U2OS cells were precipitated with immobilized Abs specific to UFL1_301-389_ (**a, e, i, m, q**), C53 (**b, f, j, n, r**), γ-tubulin (**c, g, k, o, s**), or GCP2 (**d, h, l, p, t**). The blots were probed with Abs to UFL1, C53, γ-tubulin (γ-Tb), GCP2, or GCP4. Load *(lane 1),* immobilized Abs not incubated with cell extracts *(lane 2),* precipitated proteins *(lane 3),* and carriers without Abs and incubated with cell extracts *(lane 4).* (**B**) The size distribution of proteins extracted from the membranous fraction of U2OS cells fractionated on the Superose 6 column. The blots of the collected fractions were probed with Abs to UFL1, C53, γ – tubulin (γ-Tb), GCP2, and actin. The calibration standards (in kDa) are indicated on the top. The numbers at the bottom denote individual fractions. (**C**) Pooled fractions (Nos. 15-17) from fractionation (panel **B**) were precipitated with Ab to γ-tubulin. The blots were probed with Abs to UFL1, C53, and γ-tubulin (γ-Tb). Load *(lane 1*), immobilized Ab not incubated with cell extracts (*lane 2*), precipitated proteins (*lane 3*), and the carrier without Ab and incubated with cell extracts (*lane 4*).

We separated the extract from the membranous fraction (P2) on the Superose 6B column to learn about the size of studied complexes. UFL1 was mainly distributed in high molecular weight fractions, while C53 was detected both in high molecular and low molecular weight pools. Both γ-tubulin and GCP2 were partly located in high molecular weight fractions, where γTuRCs could be present. On the other hand, the control actin was not detected in the high molecular weight pools (Fig. 1B). We confirmed the interaction of γ-tubulin with UFL1 and C53 in pooled high molecular weight fractions from the column by an immunoprecipitation experiment (Fig. 1C).

When we performed immunoprecipitation experiments with membranous fractions from glioblastoma T98G cells, we likewise detected complexes comprising UFL1, C53, and γ-tubulin (Fig. S1C), indicating that such protein interactions are not limited to U2OS cells. The formation of complexes between UFL1, C53, γ-tubulin, and GCPs was not restricted to the membranous fractions, as we found interactions between these proteins using whole-cell extracts for immunoprecipitation (Fig. S1D). Moreover, separation of wholecell extracts on the Superose 6B column resulted in partial co-distribution of UFL1, C53, γ-tubulin, and GCP2 in high molecular weight pools (Fig. S1E). Altogether, these results document that the mutually interacting UFL1 and C53 form complexes with γTuRC proteins.

### Association of exogenous UFL1 and C53 with γTuRC proteins and centrosome

To independently validate the interaction of UFL1 and C53 with γTuRC proteins, we performed immunoprecipitation experiments from cells expressing EGFP-tagged UFL1 or C53 and control cells expressing EGFP alone. The Ab to GFP co-precipitated γ-tubulin, GCP2, GCP4, GCP6, and C53 from the cells expressing EGFP-UFL1 or C53-EGFP (Fig. 2A). Reciprocal precipitation with Ab to γ-tubulin confirmed the interaction of EGFP-tagged UFL1 or EGFP-tagged C53 with γ-tubulin (Fig. 2B). On the other hand, the Ab to GFP did not co-precipitate UFL1, C53, γ-tubulin, GCP2, or GCP4 from control cells expressing EGFP alone. Similarly, Ab to γ-tubulin did not co-precipitate EGFP from the cells expressing EGFP alone (Fig. S1G). The isotype controls for the immunoprecipitation experiments with rabbit and mouse Abs are shown in Fig. S1F.

**Figure 2.**
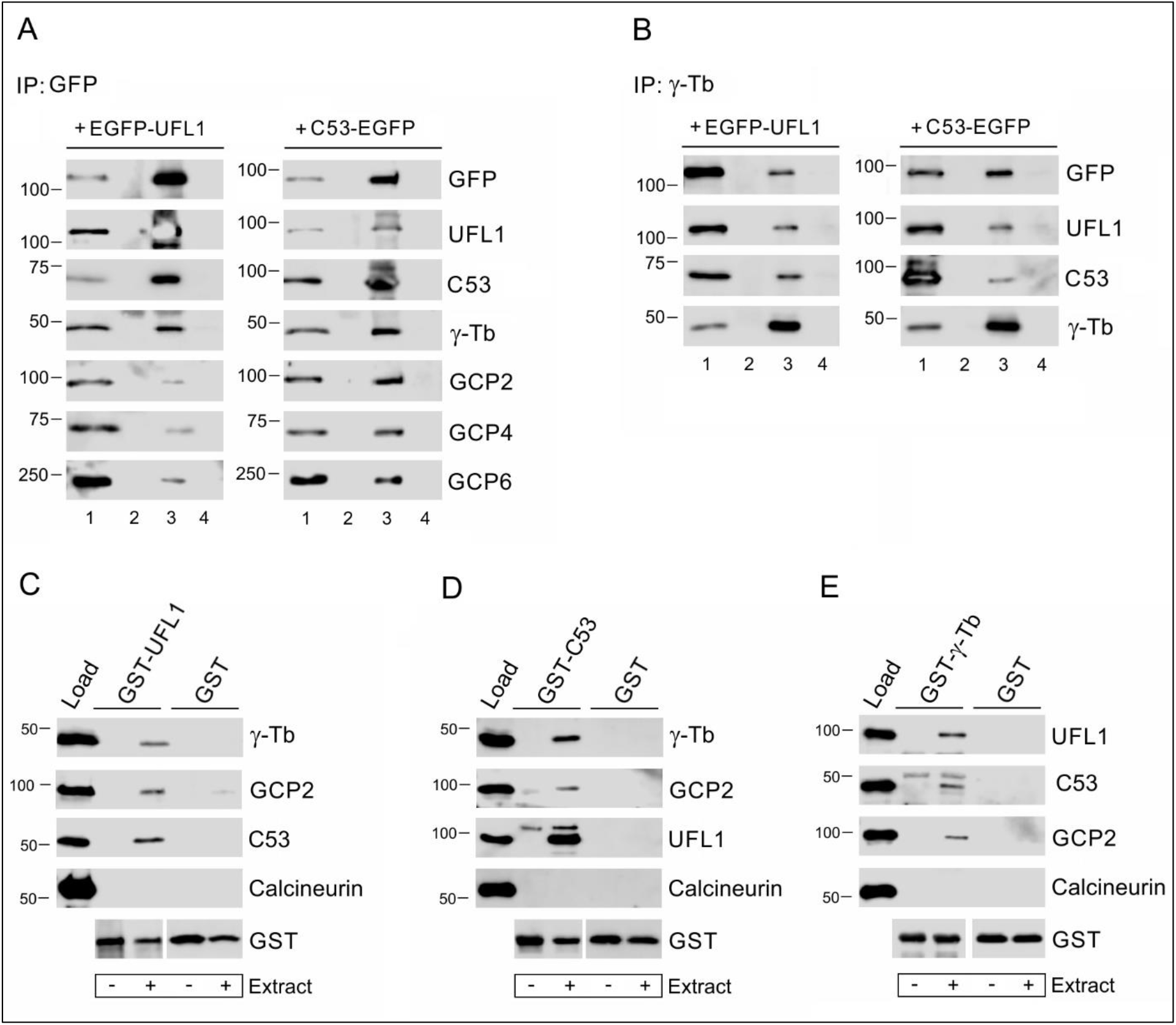
Exogenous UFL1 and C53 interact with γTuRC proteins. (**A-B**) Immunoprecipitation experiments with the whole-cell extract from U2OS cells expressing EGFP-tagged UFL1 (EGFP-UFL1) or C53 (C53-EGFP). (**A**) Precipitation with immobilized Ab to GFP. (**B**) Precipitation with immobilized Ab to γ-tubulin. The blots were probed with Abs to GFP, UFL1, C53, γ-tubulin (γ-Tb), GCP2, GCP4, or GCP6. Load *(lane 1),* immobilized Abs not incubated with cell extracts *(lane 2),* precipitated proteins *(lane 3),* and carriers without Abs and incubated with cell extracts *(lane 4).* (**C-E**) Pull-down assay with GST-tagged UFL1 (**C**), C53 (**D**), and γ-tubulin (**E**). Immobilized GST-fusion proteins (GST-UFL1, GST-C53, GST-γ-tubulin) or immobilized GST alone were incubated with whole-cell extracts from U2OS cells (Load). Blots of bound proteins were probed with Abs to γ-tubulin (γ-Tb), GCP2, UFL1 and C53, calcineurin (negative control), and GST.

To further verify the interaction of UFL1 and C53 with γTuRC proteins, we performed pull-down assays with GST-tagged C53, UFL1, or γ-tubulin. The experiments revealed that γ-tubulin and GCP2 bound to GST-UFL1 or GST-C53, but not to GST alone (Fig. 2, C and D). Similarly, UFL1 and C53 bound to GST-tagged γ-tubulin but not to GST alone (Fig. 2E). The negative control protein (calcineurin) did not bind to GST-fusion proteins, although the amounts of immobilized GST fusion proteins were comparable, as evidenced by staining with Ab to GST (Fig. 2, C-E). Collectively, these data strongly suggest that exogenous UFL1 and C53 also form complexes with γTuRC proteins.

As γTuRCs are essential for normal microtubule nucleation from centrosomes, we tested whether UFL1 or C53 localize to the centrosome. However, using immunofluorescence microscopy with a limited number of commercially available Abs to UFL1 and C53, we failed to localize these proteins on the centrosome. We, therefore, expressed TagRFP-tagged UFL1, C53, or protein tyrosine phosphatase SHP-1 (negative control) in U2OS cells. While C53-TagRFP localized to interphase centrosomes, cytosol, and nuclei (Fig. 3A, a-c), UFL1-TagRFP was found only in the cytosol (Fig. 3A, d-f), similarly to control SHP-1-TagRFP (Fig. 3A, g-i). C53-TagRFP was also clearly detected on centrosomes in living cells (Fig. 3B, a-c), in contrast to UFL1-TagRFP. These data document that although UFL1 and C53 interact with each other and form complexes with γTuRC proteins, only C53 was detected at the centrosome.

**Figure 3.**
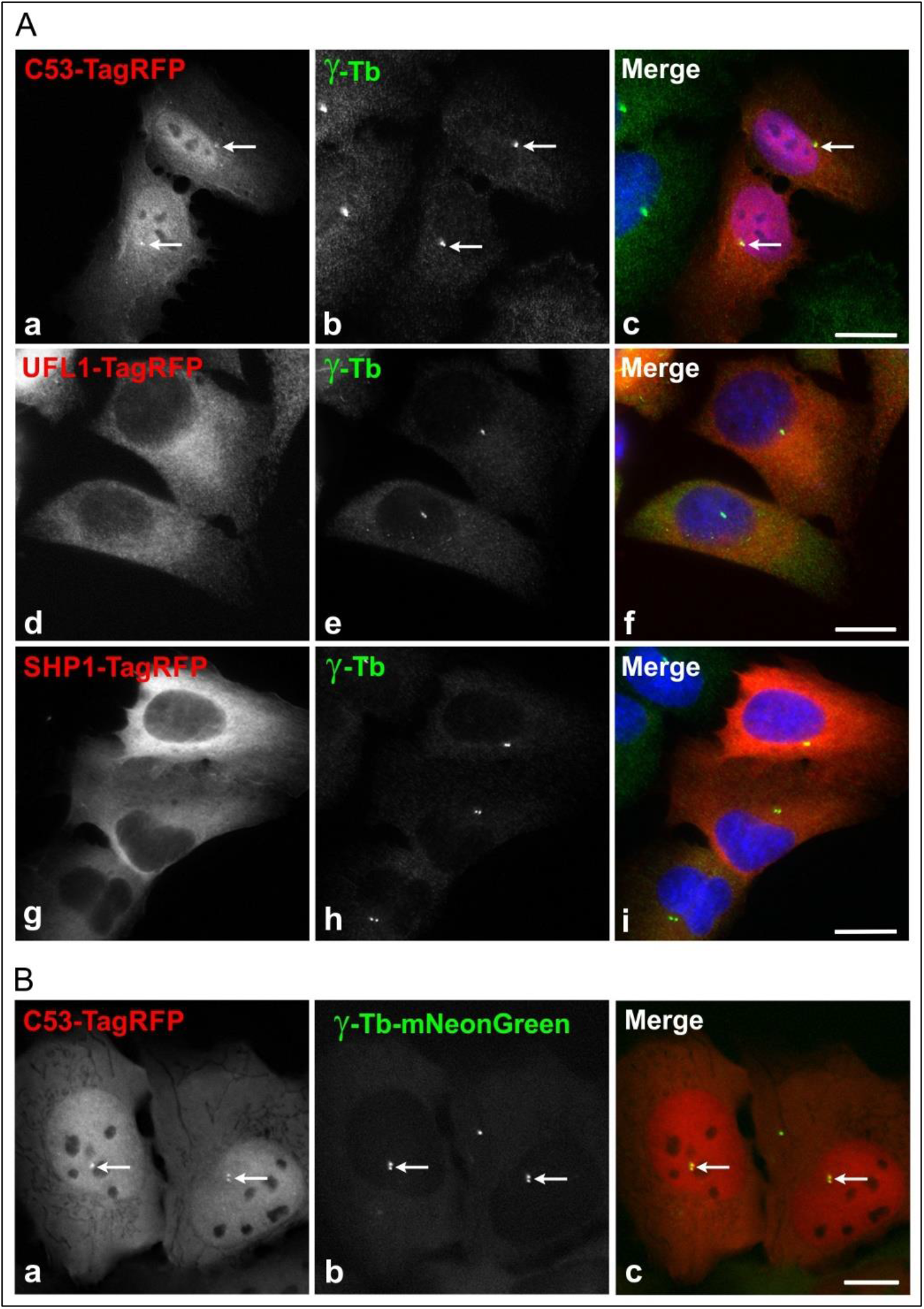
Subcellular localization of exogenous UFL1 and C53 in interphase cells. (**A**) U2OS cells expressing TagRFP-tagged proteins were fixed and stained with Ab to γ-tubulin. (**a-c**) Localization of C53-TagRFP (**a**) and γ-tubulin (**b**). Superposition of images (**c**, C53-TagRFP, red; γ-tubulin, green; DAPI, blue). (**d-f**) Localization of UFL1-TagRFP (**d**) and γ-tubulin (**e**). Superposition of images (**f**, UFL1-TagRFP, red; γ-tubulin, green; DAPI, blue). (**g-i**) Localization of negative control SHP-1-TagRFP (**g**) and γ-tubulin (**h**). Superposition of images (**i**, SHP-1-TagRFP, red; γ-tubulin, green; DAPI, blue). Arrows indicate the same positions. Fixation D/F/M. Scale bar, 20 μm. (**B**) Live cell imaging of cells expressing C53-TagRFP and γ-tubulin-mNeonGreen. (**a-c**) Localization of C53-TagRFP (**a**) and γ-tubulin-mNeonGreen (**b**). Superposition of images (**c**, C53-TagRFP, red; γ-tubulin-mNeonGreen, green; DAPI, blue). Arrows indicate the same positions. Scale bar, 10 μm.

### Preparation and characterization of cell lines lacking UFL1 or C53

To evaluate the possible effect of UFL1 and C53 on microtubule nucleation, we prepared U2OS cell lines lacking UFL1. For that, we took advantage of CRISPR/Cas 9 editing. To delete the gene region containing the canonical and alternative start codons, cells were transfected with CRISPR/Cas9 vectors (sgRNA#1, sgRNA#2, SpCas9) together with reporter plasmid pRR-hUFL1-puro for the enrichment of cells not expressing UFL1. A schematic diagram of the human *UFL1* gene with sites targeted by sgRNA#1 and sgRNA#2, enabling efficient deletion of all UFL1 isoforms, is shown in Fig. S2A. We established three independent cell lines (denoted UFL1_KO1, UFL1_KO2, and UFL1_KO3) that have deletions in the targeted region (Fig. S2B) and undetectable UFL1 in immunoblotting (Fig. S2C). When compared to control cells, a radical decrease in immunofluorescence staining was observed in UFL1_KO cells (Fig. S2D) with Ab to UFL1. If not mentioned otherwise, the following results are based on UFL1_KO1 cells (abbreviated UFL1_KO).

We used the same approach to prepare U2OS cells lacking C53. For enrichment of cells not expressing C53, we prepared reporter plasmid pRR-hCDK5RAP3-puro. A schematic diagram of the human *CDK5RAP3* gene with sites targeted by sgRNA#1 and sgRNA#2, enabling efficient deletion of all C53 isoforms, is shown in Fig. S2E. We established three independent cell lines (denoted C53_KO1, C53_KO2, and C53_KO3) that have deletions in the targeted region (Fig. S2F) and undetectable C53 in immunoblotting (Fig. S2G). When compared to control cells, a substantial decrease in immunofluorescence staining was observed in C53_KO cells (Fig. S2H). If not mentioned otherwise, the following results are based on C53_KO1 cells (abbreviated C53_KO).

To assess the effect of UFL1 or C53 deletion on cell division, we determined cell growth in control and UFL1_KO or C53_KO cells. When compared to control cells, the number of viable cells decreased both in UFL1-KO and C53_KO, but proliferation was hampered more in UFL1_KO cells (Fig. 4A). Quantitative immunoblot analysis showed that the deletion of UFL1 resulted in a substantial reduction of C53 and DDRGK1. On the other hand, the deletion of C53 only resulted in a moderate decrease in UFL1 and DDRGK1. In cells lacking UFL1, the amount of C53 and DDRGK1 dropped to ~10% and ~40% of the wildtype level, respectively. In cells lacking C53, the amount of UFL1 and DDRGK1 decreased to ~75% and ~80% of the wild-type level, respectively (Fig. 4B). The deletion of UFL1 or C53 did not affect the expression of γ-tubulin or GCP2. Changes in cell proliferation were also documented by quantitative immunoblot analysis with Abs to cyclin B1 and mitotic p-Histone H3. Lower amounts of cyclin B1 and p-Histone H3 were detected in UFL1_KO cells when compared to controls. A similar tendency, but less prominent, was observed in the case of C53_KO cells (Fig. 4C).

**Figure 4.**
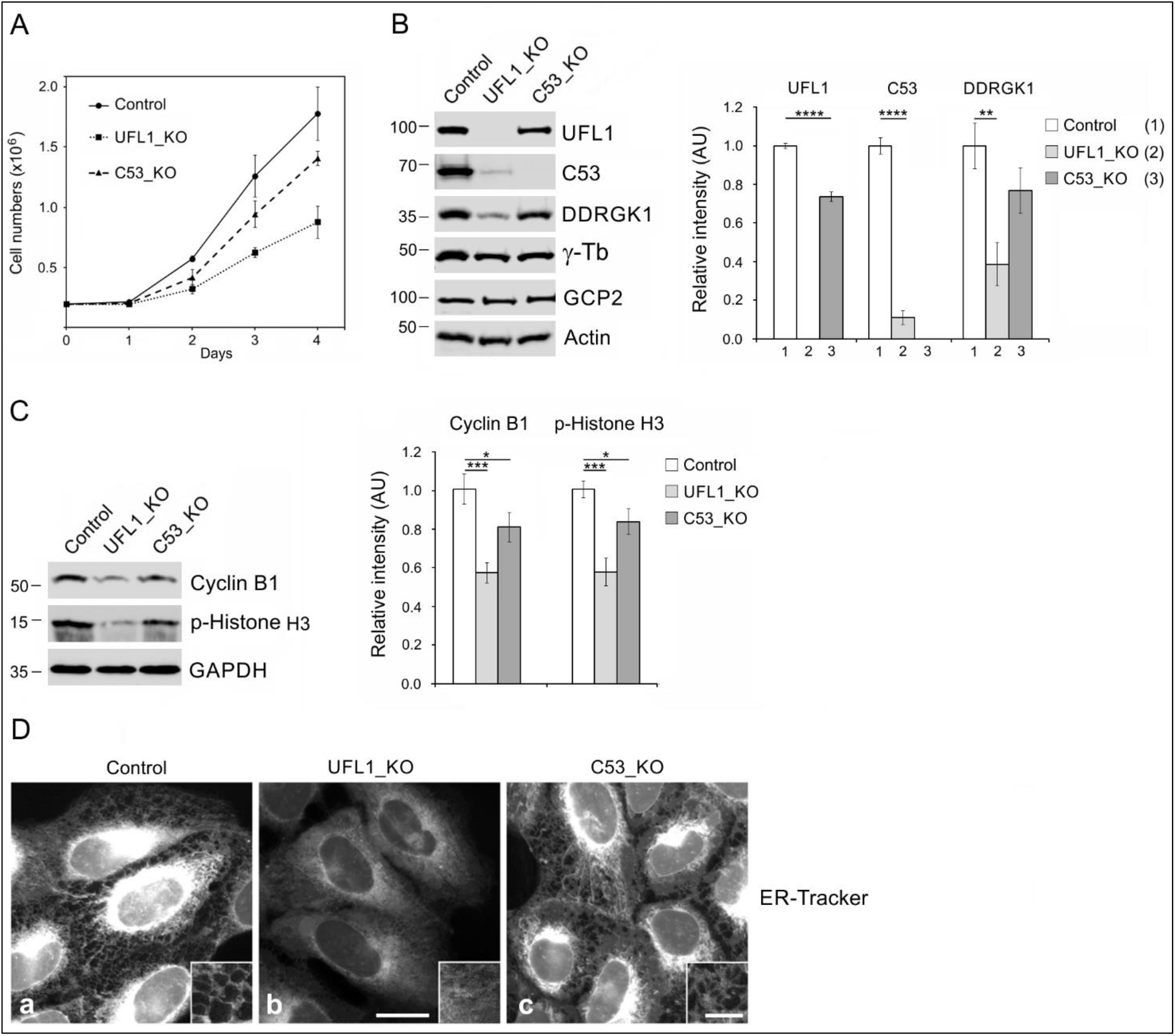
Characterization of cells lacking UFL1 or C53. (**A**) Growth curves in control, UFL1-deficient (UFL1_KO), or C53-deficient (C53_KO) U2OS cells. A total of 2 x 10^5^ cells was plated for each cell line. The values indicate mean ± SD (n=2). (**B**) Changes in the expression of UFL1, C53, and DDRGK1. The blots from whole-cell lysates were probed with Abs to UFL1, C53, DDRGK1, γ-tubulin (γ-Tb), GCP2, and actin (loading control). Densitometric quantification of immunoblots is shown on the right. Relative intensities of corresponding proteins normalized to control cells and the amount of actin in individual samples. Values indicate mean ± SD (n=3). (**C**) Changes in the expression of cyclin B1 and p-histone H3. The blots from whole-cell lysates were probed with Abs to Cyclin B1, p-Histone H3, and GAPDH (loading control). Densitometric quantification of immunoblots is shown on the right. Relative intensities of corresponding proteins normalized to control cells and the amount of GAPDH in individual samples. Values indicate mean ± SD (n=3). One-way ANOVA with Sidak’s multiple comparisons test was performed to determine statistical significance. *, p < 0.05, **, p < 0.01, ***, p < 0.001, ****, p < 0.0001. (**D**). Distribution of ER in control (**a**), UFL1_KO **(b**), and C53_KO (**c**) cells visualized by ER-Tracker in living cells. A higher magnification view of the cell periphery is shown in the image inset (**a-c**). The images (**a, b, c**) were collected and processed in the same manner. Scale bars, 20 μm (**b**) and 5 μm (inset in **c**).

Experiments on knockout mouse models revealed that the deletion of UFL1 (Li et al., 2018; Zhang et al., 2015) or C53 (Y ang et al., 2019) triggered ER stress and activated the unfolded protein response (UPR). One of the characteristic features of stressed cells is the expansion of the ER network (Sriburi et al., 2004). To characterize the prepared cell lines with respect to ER stress, we visualized ER in living cells by cellpermeant ER-Tracker. Substantial changes in ER distribution were observed both in UFL1_KO and C53_KO cells. In control cells, the staining intensity of ER-Tracker cumulated in the perinuclear region. The ER tubules at the cell periphery were sparse (Fig. 4D, a). In UFL1_KO cells, ER-Tracker staining intensity was more evenly distributed throughout the cells, suggesting the expansion of the ER network (Fig. 4D, b). The ER expansion in the C53_KO cell was less prominent than in UFL1_KO cells (Fig. 4D, c).

Collectively these findings point to the vital role of UFL1 and C53 in cell proliferation and ER homeostasis. In good agreement with previous studies (Kwon et al., 2010; Wu et al., 2010), UFL1 effectively regulates the C53 protein level.

### UFL1 or C53 deficiency increase centrosomal microtubule nucleation

To unravel the effect of UFL1 and C53 deficiency on microtubule nucleation, we followed microtubule regrowth from U2OS centrosomes, which represent the major nucleation centers in interphase cells. Microtubule regrowth in nocodazole-washout experiments in control, UFL1_KO, and C53_KO cells were performed as previously described (Sulimenko et al., 2015). The extent of microtubule regrowth could be modulated by mechanisms regulating either microtubule nucleation or microtubule dynamics. It was reported previously that microtubule dynamics is regulated in the cell periphery and that a delay in microtubule regrowth is associated with defects in microtubule nucleation (Colello et al., 2010). We measured α-tubulin and γ-tubulin immunofluorescence signals in a 2.0 μm ROI, 2.0 min after nocodazole-washout in UFL1- or C53-deficient cells and controls. When compared with control cells, an increase in microtubule regrowth was observed both in UFL1_KO1 (Fig. 5A) and C53_KO1 (Fig. 5D) cells. Quantification of γ-tubulin immunofluorescence revealed that the amount of γ-tubulin in centrosomes increased in both UFL1_KO1 (Fig. 5B) and C53_KO1 (Fig. 5E) cells. Typical staining of α-tubulin and γ-tubulin in control, UFL1_KO1, and C53_KO1 cells is shown in Fig. 5C and Fig. 5F. When microtubule regrowth experiments were performed with the other deficient cell lines (UFL1_KO2, UFL1_KO3 or C53_KO2, C53_KO3), similar results were obtained. While the amount of centrosomal γ-tubulin increased in the cells lacking UFL1 or C53, the amount of centrosomal pericentrin was not affected (Fig. S3, A-D).

**Figure 5.**
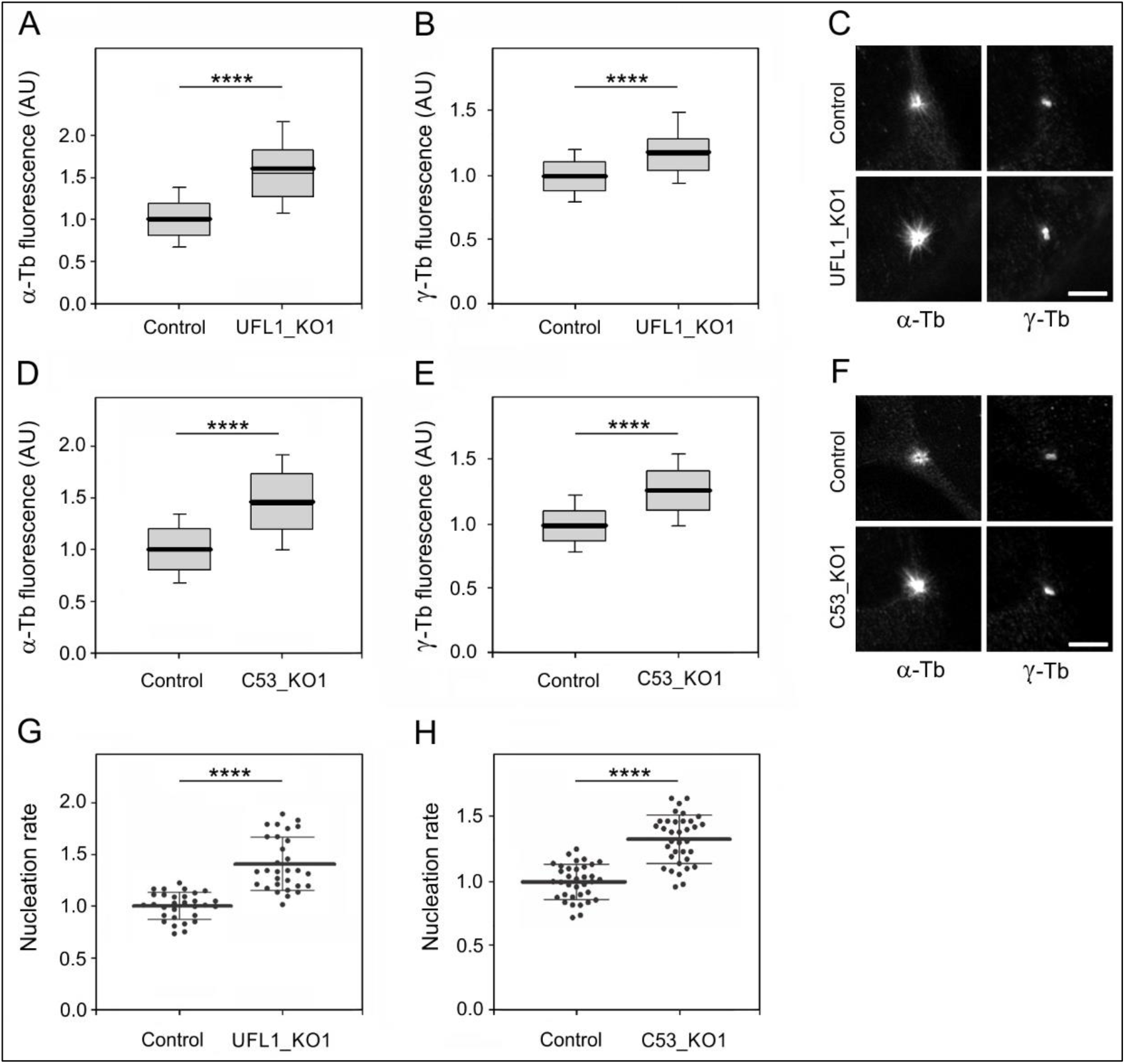
Deletion of UFL1 or C53 increases centrosomal microtubule nucleation. Centrosomal microtubule nucleation was evaluated by quantification of microtubule regrowth in fixed cells (**A-F**) and by measuring the microtubule nucleation rate in living cells (**G-H**). (**A-B, D-E**) The distribution of α-tubulin or γ-tubulin fluorescence intensities (arbitrary units [AU]) in 2-μm ROI at 2.0 min of microtubule regrowth are shown as box plots (three independent experiments, > 58 cells counted for each experimental condition). (**A-B**) Box plot of α-tubulin (**A**) and γ-tubulin **(B**) fluorescence intensities in UFL1_KO1 cells (n=239) relative to control cells (n=237). (**D-E**) Box plot of α-tubulin (**D**) and γ-tubulin (**E**) fluorescence intensities in C53_KO1 cells (n=257) relative to control cells (n=274). The bold and thin lines within the box represent mean and median (the 50th percentile), respectively. The bottom and top of the box represent the 25th and 75th percentiles. Whiskers below and above the box indicate the 10th and 90th percentiles. (**C, F**) Labeling of α-tubulin and γ-tubulin in the microtubule regrowth experiment in the control and UFL1_KO cells (**C**) or the control and C53_KO1 cells (**F**). Cells were fixed (F/Tx/M) at 2.0 min of microtubule regrowth. The pairs of images (α-Tb), (γ-Tb) were collected and processed in the same manner. Scale bars, 5 μm. (**G**) Microtubule nucleation rate (EB3 comets/min) in UFL1_KO1 cells relative to controls. Three independent experiments (at least 10 cells counted in each experiment). Control (n=30), UFL1_KO1 (n=30). The bold and thin lines within dot plot represent mean ± SD. (**H**) Microtubule nucleation rate (EB3 comets/min) in C53_KO1 cells relative to controls. Three independent experiments (at least 10 cells counted in each experiment). Control (n=35), C53_KO1 (n=35). The bold and thin lines within dot plot represent mean ± SD. Two-tailed, unpaired Student’s *t* test was performed to determine statistical significance. ****, p < 0.0001.

To independently evaluate the role of UFL1 and C53 in microtubule nucleation, we performed time-lapse imaging in cells expressing mNeonGreen-tagged microtubule end-binding protein 3 (EB3), decorating plus ends of the growing microtubules (Akhmanova and Steinmetz, 2008), and counted the number of EB3 comets leaving the centrosomes per unit time (nucleation rate) (Černohorská et al., 2016; Colello et al., 2010). When compared to control cells, the nucleation rate increased in cells lacking UFL1 (Fig. 5G). A similar effect was observed after deletion of C53 (Fig. 5H). A comparison of a single frame or 10-frame projections from control and UFL1_KO1 or C53_KO1 cells is shown in Fig. S3E and Fig. S3F. These live-imaging data corresponded to the results obtained by measuring the α-tubulin signal during the microtubule regrowth experiment.

To verify the specificity of observed changes in microtubule nucleation, we performed rescue experiments by expressing UFL1-TagRFP or TagRFP alone in UFL1_KO cells. The introduction of UFL1-TagRFP into UFL1-deficient cells restored UFL1 expression and enhanced the expression of C53 (Fig. 6A). While the introduction of UFL1-TagRFP into UFL1_KO cells decreased the microtubule regrowth to that in control cells, the expression of TagRFP failed to do so (Fig. 6B). Correspondingly, the amount of centrosomal γ-tubulin decreased after the introduction of UFL1-TagRFP into UFL1-KO cells, whereas it remained elevated in UFL1_KO cells expressing TagRFP alone (Fig. 6C). A similar set of phenotypic rescue experiments was performed in C53_KO cells. C53-TagRFP efficiently restored the C53 level in deficient cells (Fig. 6D), and microtubule regrowth was restored to that in control cells (Fig. 6E). Also, the amount of γ-tubulin in centrosomes decreased after introducing C53-TagRFP to deficient cells (Fig. 6F).

**Figure 6.**
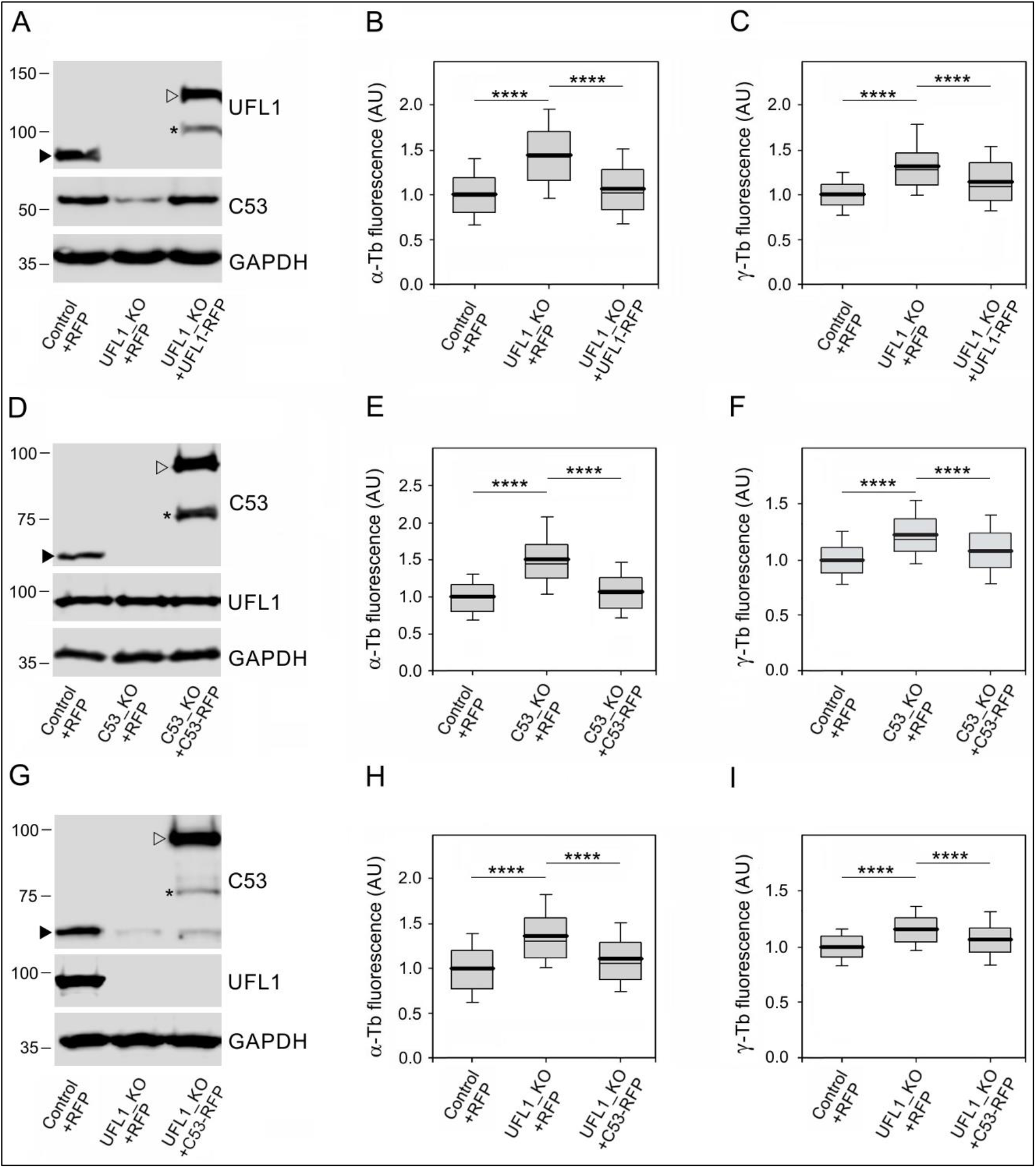
C53 is sufficient to attenuate centrosomal microtubule nucleation in *UFL1* knockout cells. (**A**) Immunoblot analysis of UFL1 and C53 in whole-cell lysates from control cells expressing TagRFP (Control+RFP), UFL1_KO cells expressing TagRFP (UFL1_KO+RFP), and UFL1_KO cells rescued by UFL1-TagRFP (UFL1_KO+UFL1-RFP). Blots were probed with Abs to UFL1, C53, and GAPDH (loading control). Black and empty arrowheads and asterisk denote, respectively, endogenous UFL1, UFL1-TagRFP, and its fragment. (**B-C**) The distributions of α-tubulin or γ-tubulin fluorescence intensities (arbitrary units [AU]) in 2-μm ROI at 2.0 min of microtubule regrowth are shown as box plots (four independent experiments, > 30 cells counted for each experimental condition). Box plot of α-tubulin (**B**) and γ-tubulin **(C**) fluorescence intensities in UFL1_KO+RFP (n=181) and UFL1_KO+UFL1-RFP cells (n=298) relative to control cells (Control+RFP, n=267). (**D**) Immunoblot analysis of C53 and UFL1 in whole-cell lysates from control cells expressing TagRFP (Control+RFP), C53_KO cells expressing TagRFP (C53_KO+RFP), and C53_KO cells rescued by C53-TagRFP (C53_KO+C53-RFP). Blots were probed with Abs to C53, UFL1, and GAPDH (loading control). Black and empty arrowheads and asterisk denote, respectively, endogenous C53, C53-TagRFP, and its fragment. (**E-F**) The distributions of α-tubulin or γ-tubulin fluorescence intensities (arbitrary units [AU]) in 2-μm ROI at 2.0 min of microtubule regrowth are shown as box plots (three independent experiments, > 30 cells counted for each experimental condition). Box plot of α-tubulin (**E**) and γ-tubulin (**F**) fluorescence intensities in C53_KO+RFP (n=191) and C53_KO+C53-RFP cells (n=248) relative to control cells (Control+RFP, n=179). (**G**) Immunoblot analysis of C53 and UFL1 in whole-cell lysates from control cells expressing TagRFP (Control+RFP), UFL1_KO cells expressing TagRFP (UFL1_KO+RFP), and UFL1_KO cells rescued by C53-TagRFP (UFL1_KO+C53-RFP). Blots were probed with Abs to C53, UFL1, and GAPDH (loading control). Black and empty arrowheads and asterisk denote, respectively, endogenous C53, C53-TagRFP, and its fragment. (**H-I**) The distributions of α-tubulin or γ-tubulin fluorescence intensities (arbitrary units [AU]) in 2-μm ROI at 2.0 min of microtubule regrowth are shown as box plots (three independent experiments, > 37 cells counted for each experimental condition). Box plot of α-tubulin (**H**) and γ-tubulin (**I**) fluorescence intensities in UFL1_KO+RFP (n=133) and UFL1_KO+C53-RFP (n=152) relative to control cells (Control+RFP, n=174). (**B, C, E, F, H, I**) Bold and thin lines within the box represent mean and median (the 50th percentile), respectively. The bottom and top of the box represent the 25th and 75th percentiles. Whiskers below and above the box indicate the 10th and 90th percentiles. One-way ANOVA with Sidak’s multiple comparisons test was performed to determine statistical significance. ****, p < 0.0001.

To test whether C53 can rescue increased microtubule nucleation phenotype in UFL1_KO cells, C53-TagRFP or TagRFP alone were expressed in UFL1_KO cells. C53-TagRFP was efficiently expressed in UFL1_KO cells that are characterized by a low amount of C53 (Fig. 4B and Fig. 6G). In UFL1_KO cells expressing C53-TagRFP, both microtubule regrowth (Fig. 6H) and the amount of centrosomal γ-tubulin (Fig. 6I) decreased to the levels in control cells. These data highlight C53 as a novel regulator of centrosomal microtubule nucleation.

Altogether, these data indicate that both UFL1 and C53 negatively regulate microtubule nucleation from the interphase centrosome by influencing centrosomal γ-tubulin/γTuRCs levels. The modulation of microtubule nucleation by UFL1 likely occurs by reducing the amount of centrosomal C53.

### Enhancement of microtubule nucleation in cells under ER stress

As live-cell imaging of ER in control, UFL1_KO, and C53_KO cells indicated generation of ER stress in cells lacking UFL1 and C53 (Fig. 4D), we evaluated changes in the expression and cellular distribution of chaperone calnexin and protein disulfide-isomerase (PDI), well-established markers of UPR, facilitating protein folding (Chakrabarti et al., 2011). Densitometric analysis of immunoblotting experiments revealed a significantly increased expression of calnexin and PDI in UFL1_KO cells. These UPR markers also increased in C53_KO cells, albeit PDI only moderately (Fig. 7A). Immunofluorescence microscopy verified increased expression of calnexin (Fig. 7B, e and i) and PDI (Fig. S4A, e and i) in cells lacking UFL1 or C53. When compared with control cells, immunostaining for calnexin spread to the cell periphery delineated by ends of microtubules both in UFL1_KO and C53_KO cells, as shown in magnified regions (Fig. 7B, g and k). Similarly, the PDI immunostaining also expanded to the cell periphery in cells lacking UFL1 or C53 (Fig. S4A, g and k). To evaluate whether C53 can rescue calnexin spreading to the cell periphery in UFL1_KO cells, C53-TagRFP or TagRFP alone were expressed in UFL1_KO cells. In UFL1_KO cells expressing C53-TagRFP, the distribution of calnexin in the cell periphery approached that in control cells (Fig. 7C, a-c). Staining for TagRFP delineated cell periphery (Fig. 7C, d-f). Densitometric analysis of immunoblots revealed a partial decrease in calnexin protein level in UFL1_KO cells expressing C53-TagRFP (Fig. S4B), indicating that C53 cannot completely rescue significant ER stress induced in UFL1-KO cells lacking the ufmylation capability.

**Figure 7.**
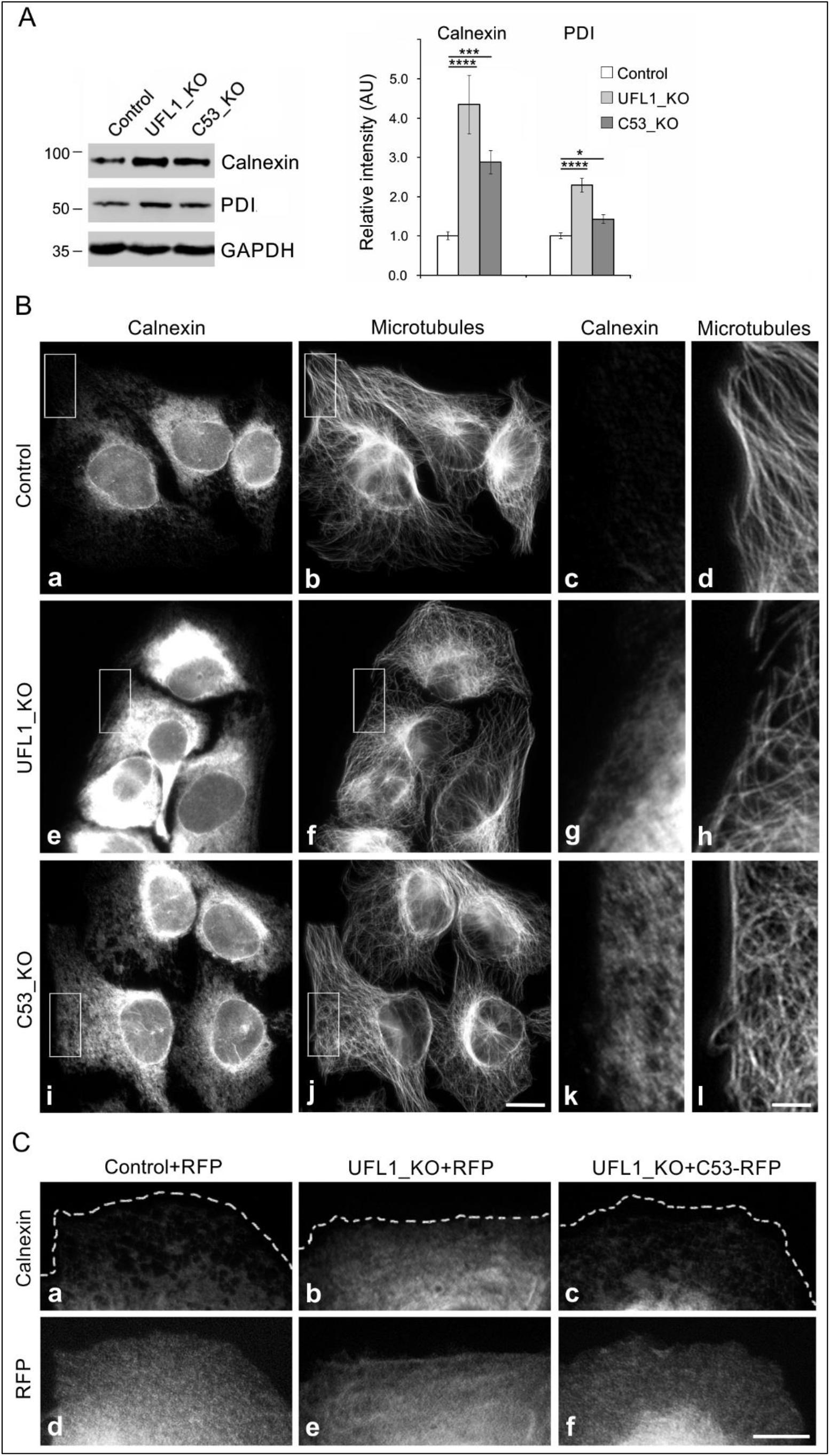
Deletion of UFL1 or C53 induces UPR and leads to relocation of calnexin to the cell periphery. (**A**) Immunoblot analysis of calnexin and PDI in U2OS cells lacking UFL1 or C53. The blots from whole-cell lysates were probed with Abs to calnexin, PDI, and GAPDH (loading control). Densitometric quantification of immunoblots is shown on the right. Relative intensities of corresponding proteins normalized to control cells and the amount of GAPDH in individual samples. Values indicate mean ± SD (n=4 for calnexin; n=3 for PDI). One-way ANOVA with Sidak’s multiple comparisons test was performed to determine statistical significance, *, p < 0.05, ***, p < 0.001, ****, p < 0.0001. (**B**) Immunofluorescence microscopy. (**a-d**) Control U2OS cells, (**e-h**) UFL1-deficient cells (UFL1_KO) and (**i-l**) C53-deficient cells (C53_KO). Cells were fixed and double-labeled for calnexin (**a, e, i**) and β-tubulin (**b, f, j**; Microtubules). Higher magnification views of the regions delimited by rectangles are shown on the right of images from control (**c-d**), UFL1_KO (**g-h**), and C53_KO (**k-l**) cells. The images (**a, e, i**) and (**c, g, k**) were collected and processed in the same manner. Fixation F/Tx. Scale bars, 20 μm (**j**), and 5 μm (**l**). (**C**) Calnexin localization in a phenotypic rescue experiment. (**a, d**) Control cells expressing TagRFP (Control+RFP), (**b, e**) UFL1_KO cells expressing TagRFP (UFL1_KO+RFP), and (**c, f**) UFL1_KO cells rescued by C53-TagRFP (UFL1_KO+C53-RFP). Cells were fixed and stained for calnexin (**a-c**). The images (**a, b, c**) were collected and processed in the same manner. Fixation F/Tx. Scale bar, 10 μm.

Microtubules are known to regulate ER homeostasis, with ER dynamics tightly linked to the dynamics of microtubules. This is particularly important during ER stress, as the expansion of the ER is one of the relief mechanisms (Schuck et al., 2009). To evaluate whether the expansion of ER in stressed cells could be potentially promoted by *de novo* microtubule nucleation, we pretreated U2OS cells with tunicamycin, a potent inhibitor of protein glycosylation and ER stress activator (Ding et al., 2007). Immunofluorescence microscopy in tunicamycin-treated cells showed expansion of ER, as documented by staining of live cells with ER-Tracker which was more evenly distributed throughout treated-cells than in controls (Fig. 8A, a and b), and by staining fixed cells with Abs to calnexin (Fig. 8A, c and d) and PDI (Fig. 8A, e and f). Moreover, DNA damage-inducible transcript 3 (DDIT3), representing another UPR marker (Chakrabarti et al., 2011), accumulated in nuclei after tunicamycin treatment (Fig. 8A, g and h). Increased expression of calnexin, PDI, and DDIT3 after tunicamycin treatment was confirmed by immunoblotting (Fig. 8B). Changes in the protein level corresponded to changes in the transcript level (Fig. S4C). Interestingly, we detected increased microtubule regrowth (Fig. 8C) and centrosomal γ-tubulin accumulation (Fig. 8D) in tunicamycin-treated cells. Correspondingly, the nucleation rate also increased in treated cells (Fig. 8E and Fig. 8F). When cells expressing C53-TagRFP were pretreated with tunicamycin, fluorescence microscopy revealed that the overall C53-TagRFP signal decreased, and its association with the centrosome was suppressed (Fig. 9A). Immunoblot analysis showed that while the amount of C53-TagRFP diminished in tunicamycin-treated cells, protein levels of UFL1, endogenous C53 or γ-tubulin were similar in control and drug-treated cells (Fig. 9B). No changes in fluorescence staining intensity were observed in cells expressing TagRFP alone and treated or not with tunicamycin (Fig. 9C). Immunoblot analysis also did not reveal changes in the amount of TagRFP (Fig. 9D), ruling out the possibility that tunicamycin affects expression of tagged proteins. Interestingly, lower amount of C53 associated with P1 fraction, comprising centrosomes and nuclei with connected membranes, in tunicamycin-treated cells. On the other hand, a comparable amount of pericentrin was found in P1 fractions of control and drug-treated cells (Fig. 9E). Obtained data indicate that tunicamycin induced subcellular redistribution of C53 away from the centrosome.

**Figure 8.**
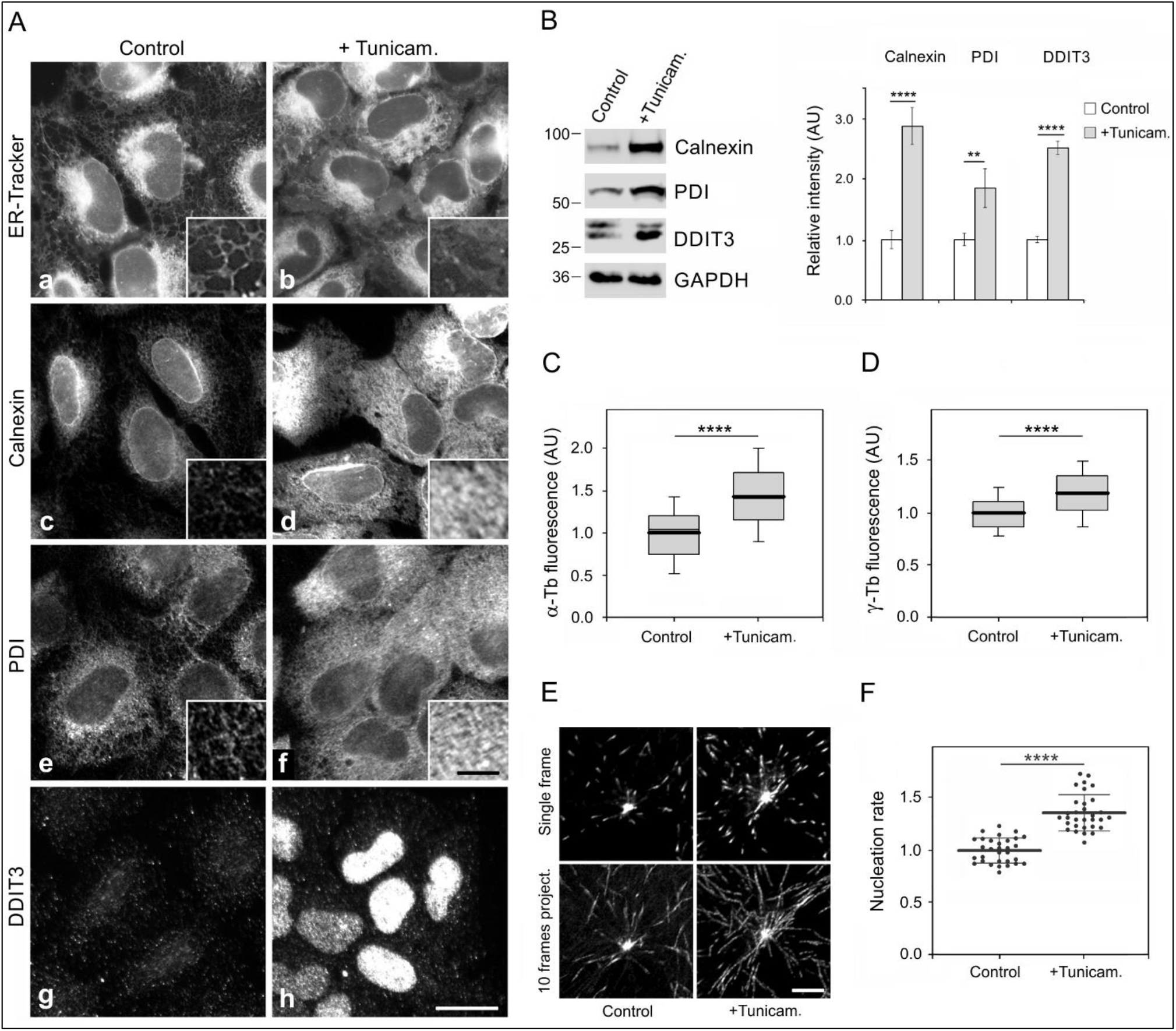
Generation of ER stress by tunicamycin induces UPR and increases centrosomal microtubule nucleation. U2OS cells were treated with 1 μg/ml tunicamycin (+Tunicam.) or DMSO carrier (Control) for 24 h. (**A**). Effect of tunicamycin on expression and subcellular distribution of calnexin or PDI, and ER stress-induced transcription factor DDIT3. (**a-b**) Visualization of ER in living cells by ER-Tracker. (**c-h**) Fixed cells stained for calnexin (**c-d**), PDI (**e-f**), and DDIT3 (**g-h**). A higher magnification view of the cell periphery is shown in the image insets (**a-f**). Pairs of images (**a-b**), (**c-d**), (**e-f**), and (**g-h**) were collected and processed in the same manner. Fixation F/Tx. Scale bars, 20 μm (**h**) and 5 μm (inset in **f**). (**B**) Immunoblot analysis of untreated and tunicamycin-treated cells. The blots from whole-cell lysates were probed with Abs to calnexin, PDI, DDIT3, and GAPDH (loading control). Densitometric quantification of immunoblots is shown on the right. Relative intensities of corresponding proteins normalized to control cells and the amount of GAPDH in individual samples. Values indicate mean ± SD (n=5 for calnexin; n=4 for PDI; n=3 for DDIT3). (**C-D**) The distributions of α-tubulin or γ-tubulin fluorescence intensities (arbitrary units [AU]) in 2-μm ROIs at 3.0 min of microtubule regrowth in control and tunicamycin-treated cells are shown as box plots (four independent experiments, > 27 cells counted for each experimental condition). Box plot of α-tubulin (**C**) and γ-tubulin (**D**) fluorescence intensities in tunicamycin-treated cells (n=234) relative to control cells (n=181). The bold and thin lines within the box represent mean and median (the 50th percentile), respectively. The bottom and top of the box represent the 25th and 75th percentiles. Whiskers below and above the box indicate the 10th and 90th percentiles. (**E**) Time-lapse imaging of control and tunicamycin-treated cells expressing EB3-mNeonGreen. Still images of EB3 (Single frame) and tracks of EB3 comets over 10 s (10 frames project.). Scale bar, 5 μm. (**F**) Microtubule nucleation rate (EB3 comets/min) in tunicamycin-treated cells relative to control cells. Three independent experiments (at least 9 cells counted in each experiment). Control (n=31), tunicamycin-treated cells (n=31). The bold and thin lines within the dot plot represent mean ± SD. Two-tailed, unpaired Student’s *t* test was performed to determine statistical significance **, p < 0.01; **** p < 0.0001.

**Figure 9.**
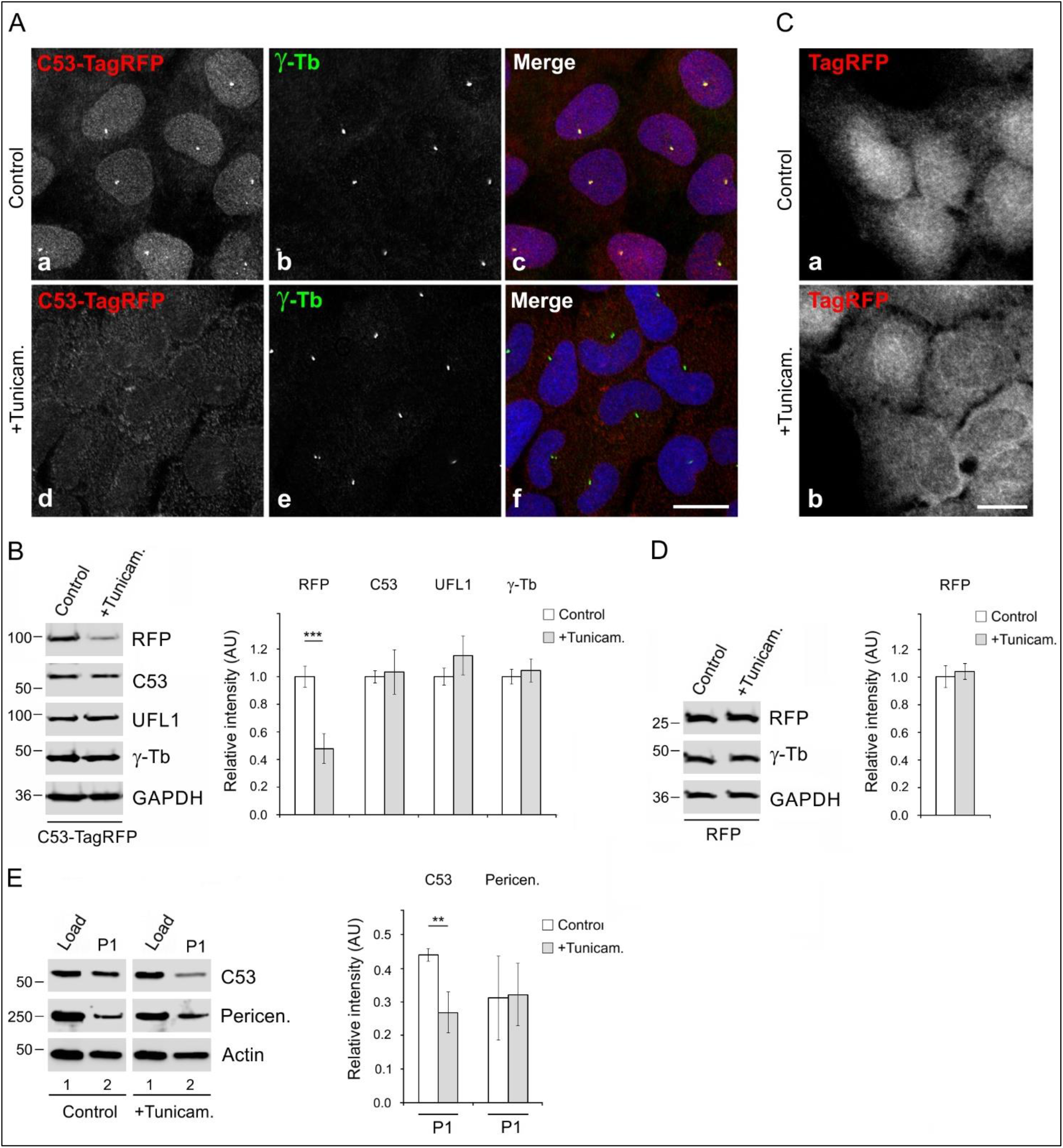
Tunicamycin affects the distribution and protein level of C53-TagRFP. U2OS cells were treated with 1 μg/ml tunicamycin (+Tunicam.) or DMSO carrier (Control) for 24 h. (**A-B**) Cells expressing C53-TagRFP. (**A**) Immunofluorescence microscopy of cells fixed and stained with Ab to γ-tubulin. (**a-c**) Control cells. C53-TagRFP (**a**), γ-tubulin (**b**), superposition of images (**c**, C53-TagRFP, red; γ-tubulin, green; DAPI, blue). (**d-f**) Tunicamycin-treated cells. C53-TagRFP (**d**), γ-tubulin (**e**), superposition of images (**f**, C53-TagRFP, red; γ-tubulin, green; DAPI, blue). Images (**a, d**) and (**b, e**) were collected and processed in the exactly same manner. Fixation Tx/F/M. Scale bar, 20 μm. (**B**) Immunoblot analysis of whole-cell lysates with Abs to RFP, C53, UFL1, γ-tubulin (γ-Tb), and GAPDH (loading control). Densitometric quantification of immunoblots is shown on the right. Relative intensities of corresponding proteins normalized to control cells and the amount of GAPDH in individual samples. Values indicate mean ± SD (n=4). (**C-D**) Cells expressing TagRFP. (**C**) Immunofluorescence microscopy of fixed control cells (**a**) and tunicamycin-treated cells (**b**). Images (**a, b**) were collected and processed in exactly the same manner. Fixation F/Tx. Scale bar, 20 μm. (**D**) Immunoblot analysis of whole-cell lysates with Abs to RFP, γ-tubulin (γ-Tb), and GAPDH (loading control). Densitometric quantification of immunoblots is shown on the right. Relative intensities of corresponding proteins normalized to control cells and the amount of GAPDH in individual samples. Values indicate mean ± SD (n=3). (**E**) Distribution of proteins in fractions after differential centrifugation of the cell homogenate. Cell fractions were prepared as described in the Materials and Methods section. Cell homogenate *(lane 1),* pellet P1 *(lane 2).* Immunoblot analysis with Abs to C53, pericentrin, and actin (loading control). Densitometric quantification of immunoblots is shown on the right. Intensities of corresponding proteins in P1 normalized to loads (relative intensity 1.0). Values indicate mean ± SD (n=4). Two-tailed, unpaired Student’s *t* test was performed to determine statistical significance. ** p < 0.01; ***, p < 0.001.

Collectively taken, these results suggest that also during pharmacologically induced ER stress, centrosomal microtubule nucleation is enhanced.

## Discussion

The ER distribution is dependent on microtubules. Although ER expansion is characteristic of cells under ER stress (Smith and Wilkinson, 2017), the molecular mechanisms of microtubule regulation in this process are not fully understood. UFL1 and its adaptor C53 are essential for ER homeostasis (Gerakis et al., 2019), and their deletions generate ER stress (Y ang et al., 2019). To evaluate the role of UFL1 and C53 in microtubule organization, we made use of well-adherent U2OS cells suitable for analysis of microtubule nucleation from interphase centrosomes. We report on UFL1 and C53 association with γTuRC proteins and their involvement in the regulation of centrosomal microtubule nucleation. Increased microtubule nucleation in cells under ER stress could facilitate ER expansion to the cell periphery.

### Interaction of UFL1 and C53 with γTuRC proteins

Several lines of evidence support the conclusion that UFL1 and C53 (UFL1/C53) associate with γTuRC proteins. First, UFL1 was identified by MALDI/MS fingerprinting analysis after immunoprecipitation with anti-peptide mAb to γ-tubulin and elution of bound proteins with the immunizing peptide. Second, reciprocal immunoprecipitations confirmed the formation of complexes containing UFL1, C53, γ-tubulin, and GCPs. Third, separation of extracts by gel filtration revealed co-distribution of UFL1, C53, γ-tubulin, and GCP2 in high molecular weight fractions, in which both UFL1 and C53 co-precipitated with γ-tubulin. Fourth, γ-tubulin and GCPs associated with EGFP-tagged UFL1 or C53. Finally, γ-tubulin and GCP2 bound to GST-tagged UFL1 or C53, and both UFL1 and C53 interacted with GST-γ-tubulin. Interaction of UFL1/C53 with γ-tubulin was observed in cells of different tissue origin. Association was found in human osteogenic sarcoma (U2OS), glioblastoma (T98G), and cervix adenocarcinoma (HeLa S3) cells. These findings suggest that multiprotein complexes containing UFL1/C53 and γTuRC proteins occur in various cell types.

UFL1 directly interacts with C53 (Kwon et al., 2010; Wu et al., 2010), which can serve as a substrate adaptor for UFL1 (Yang et al., 2019). Both UFL1 and C53 are largely ER-associated proteins (Shiwaku et al., 2010; Tatsumi et al., 2010; Wu et al., 2010). Reciprocal precipitation of these proteins from membranous fractions was therefore expected. On the other hand, the association of UFL1/C53 with membrane-bound γTuRC proteins was surprising. To our knowledge, such interaction has not been reported. We have previously shown that γ-tubulin associated with detergent-resistant membranes in mouse embryonic carcinoma P19 cells induced to neuronal differentiation (Macurek et al., 2008) and in the human brain (Dráberová et al., 2017). In this context, it should be noted that γ-tubulin is associated with vesicular structures and Golgi-derived vesicles (Chabin-Brion et al., 2001), and γ-tubulin is essential for microtubule nucleation from the Golgi membranes (Wu et al., 2016). γ-Tubulin has also been found on recycling endosomes (Hehnly and Doxsey, 2014) and mitochondrial membranes (Dráberová et al., 2017). It was reported that UFL1 regulates the mitochondrial mass (Zhang et al., 2015). Since ER and mitochondria are known to cross-talk at membrane contact sites (Lombardi and Elrod, 2017), deciphering the role of UFL1/C53 in the regulation of microtubule nucleation from membrane-bound γ-tubulin complexes warrants further investigation.

### Microtubule nucleation in cells lacking UFL1 or C53

Although UFL1/C53 interacted with γTuRC proteins, tagged fluorescent proteins revealed a centrosomal association in fixed and living cells only in the case of C53. Centrosomal localization of C53 was previously reported with antibodies (Jiang et al., 2009). We failed to localize C53 to centrosomes using a panel of commercial antibodies under various fixation conditions. The differences in immunofluorescence localization of C53 could reflect the exposure of epitopes for the used Abs. Reports on centrosomal localization of UFL1 are missing.

We prepared, using CRISPR/Cas 9 gene editing, cell lines lacking either UFL1 or C53. Cell growth inhibition, a decrease in mitotic p-Histone H3, and cyclin B1 suppression were characteristic features of UFL1_KO cells. A similar tendency, but less prominent, was also observed in the case of C53_KO cells. Decreased proliferation was described after the depletion of UFL1 by siRNA in glioma C6 cells (Shiwaku et al., 2010) and by shRNA in U2OS (Zhang et al., 2012). A decrease in proliferation and mitosis was reported in C53-deficient zebrafish embryos (Liu et al., 2011), and suppression of cyclin B1 was shown in fetal livers from C53 knockout mice (Yang et al., 2019). Our data confirm that both UFL1 and C53 participate in the regulation of the cell cycle. The absence of UFL1 resulted in a substantial reduction of C53 and DDRGK1. On the other hand, only a moderate decrease in the amount of UFL1 and DDRGK1 was observed in cells lacking C53 (Fig. 4B). It was reported that UFL1 and C53 mutually affect the stability of each other by inhibiting the ubiquitination of the other system (Kwon et al., 2010). The depletion of UFL1 rendered both C53 and DDRGK1 more susceptible to ubiquitin/proteasome system-mediated protein degradation (Wu et al., 2010). UFL1 thus plays a vital role in the regulation of the C53 protein level.

Our data demonstrate that UFL1 and C53 represent negative regulators of microtubule nucleation from interphase centrosomes. The deletion of both UFL1 and C53 increased centrosomal microtubule nucleation. Nevertheless, loss of UFL1 also leads to a strong reduction of the C53 level. The rescue of microtubule nucleation phenotype in UFL1_KO cells by C53 (Fig. 6, G-I), therefore, strongly suggests that C53 is a novel regulator of centrosomal microtubule nucleation. The regulatory role of UFL1 in microtubule nucleation could thus be indirect through modulation of the C53 amount or subcellular localization. As complexes of C53 and UFM1 have been reported (Lemaire et al., 2011; Yoo et al., 2014), it was proposed that C53 may represent an ufmylation target. However, our results from rescue experiments indicate that putative C53 ufmylation is not required for the modulation of centrosomal microtubule nucleation.

Microtubule nucleation at the centrosome occurs from γTuRCs located in the pericentriolar material (Teixidó-Travesa et al., 2012). We, therefore, examined whether UFL1 and C53 regulate microtubule nucleation by affecting the centrosomal γ-tubulin levels. Our data suggest that in wild-type cells, both proteins suppress γ-tubulin accumulation at the centrosome. On the other hand, no changes in the amount of pericentrin were detected, showing that the general pericentriolar matrix integrity is not affected by C53 or UFL1 depletion. Altogether, these data suggest that the regulatory roles of UFL1 and C53 are conveyed by γ-tubulin/γTuRC accumulation on centrosomes. Such a regulatory mechanism of microtubule nucleation is not unique only for UFL1/C53. It has been reported that androgen and Src signaling, which leads to the activation of the ERK, regulate microtubule nucleation by promoting the accumulation of γ-tubulin at the centrosome (Colello et al., 2010). Modulation of γ-tubulin accumulation in centrosomes was also shown for GIT1/βPIX signaling proteins and PAK1 kinase (Černohorská et al., 2016), and for tyrosine phosphatase SHP-1 (Klebanovych et al., 2019).

### Regulatory mechanisms by which UFL1/C53 can control microtubule nucleation

C53 binds to multiple targets but has no enzymatic domain or other well-characterized functional motifs, suggesting that it may exert its activity through interaction with other proteins. Recently, C53 was identified as a highly conserved regulator of ER-stress-induced ER-phagy (Stephani et al., 2020). Interestingly, accumulating evidence suggests that C53 is also implicated in the regulation of protein phosphorylation. It was reported that C53 directly bound to protein serine/threonine-protein phosphatase 1D (PP2Cδ) and promoted its phosphatase activity toward several C53 targets (Wamsley et al., 2017). Phosphorylation sites have been identified in building components of γTuRC. In contrast to αβ-tubulin dimers (Linhartová et al., 1992), posttranslational modification of γ-tubulin is less prominent (Vinopal et al., 2012). However, several reports have shown γ-tubulin phosphorylation (Keck et al., 2011; Kukharskyy et al., 2004; Vogel et al., 2001). Similarly, proteomic studies revealed multiple phosphorylation sites on GCPs (Teixidó-Travesa et al., 2012). Many phosphorylation sites have also been identified in various γ-TuRC tethering proteins, including NEDD1/Grip71, pericentrin, CDK5RAP2, Cnn, Mto2, and Spc110 as reviewed (Tovey and Conduit, 2018). Several of these sites have been characterized and have been shown to stimulate γTuRC assembly, recruitment, or activation (Tovey and Conduit, 2018). On the other hand, phosphorylation can also negatively regulate γTuRC recruitment and activity, as hyperphosphorylation of Mto2 in fission yeast leads to the inactivation of γTuRCs at non-spindle pole body sites (Borek et al., 2015). The activities of kinases and phosphatases have to be balanced to finely tune microtubule nucleation events during the cell cycle and in response to stress conditions. It was reported that serine/threonine-protein phosphatase 4 (PP4C) in complex with its regulatory subunits R2 and R3A associates with the centrosome, interacts with γ-tubulin and GCP2, and dephosphorylates phosphorylated γ-tubulin (Voss et al., 2013). We have recently demonstrated that tyrosine-protein phosphatase SHP-1 forms complexes with γTuRC proteins and suppresses microtubule nucleation. This indicates that different protein phosphatases might be involved in distinct signaling pathways with respect to the regulation of microtubule nucleation (Klebanovych et al., 2019). The activation of phosphatase(s) by C53 might maintain a low level of phosphorylated γTuRCs or TuRC-tethering proteins, resulting in the attenuation of microtubule nucleation.

Although the regulation of microtubule nucleation in interphase UFL1_KO cells could be explained by a low amount of C53, one cannot rule out that in mitotic cells, UFL1 affects microtubule nucleation independently of C53. It was reported that the UFM1 cascade is essential for the organization of the mitotic apparatus in *Drosophila* and alters the level of phosphorylation on tyrosine-15 of Cdk1 (pY15-Cdk1), which serves as an inhibitor of G2/M transition. In cells lacking UFL1, the level of pY15-Cdk1 was significantly reduced, indicating increased activity of Cdk1 (Yu et al., 2020). Sequential phosphorylation of NEDD1 by Cdk1 and Plk1 is required for targeting γTuRCs to centrosomes during mitosis (Zhang et al., 2009).

### Enhanced microtubule nucleation in cells under ER stress

One of the characteristic features of U2OS cells without UFL1 is the generation of ER stress, which was previously reported in mouse UFL1 knockout models for bone marrow cells (Zhang et al., 2015) and cardiomyocytes (Li et al., 2018). Similarly, U2OS cells lacking C53 are under ER stress, which was also recently reported in the mouse C53 knockout model for hepatocytes (Yang et al., 2019). When compared to control cells, where ER was unevenly distributed in the cytosol and concentrated around the nucleus, the UFL1_KO or C53_KO cells had an ER network more evenly distributed to the periphery. This was clearly evident by visualizing ER with ER-Tracker or by immunostaining for PDI or calnexin, which protein level increased in cells lacking UFL1 or C53. Expansion of ER to the cell periphery was described using staining with Ab to PDI, after partial depletion of UFL1 and C53 by RNAi (Zhang et al., 2012). Expansion of ER to the cell periphery was also clearly observable in cells pretreated with tunicamycin, a pharmacological inducer of ER stress (Ding et al., 2007).

In mammalian cells, the ER network rearrangements strongly depend on the interactions with dynamic microtubules (Terasaki et al., 1986). This is very important during ER stress, as the expansion of ER alleviates ER stress (Schuck et al., 2009; Sriburi et al., 2004). There are four distinct mechanisms of how ER can be rearranged with the help of microtubules. ER tubules can be pulled out of the existing ER membranes by associating with motor proteins and then extending along microtubules (sliding mechanism), or by attaching to the tips of growing microtubules (Waterman-Storer and Salmon, 1998). New ER tubules can also be generated by hitchhiking on organelles that are transported along microtubules by molecular motors. Finally, recent work has shown that ER tubules can be pulled by shrinking microtubule ends (Guo et al., 2018). The increased microtubule nucleation in the cells under ER stress shown in this work extends the microtubule network, which could help to expand ER membranes to the cell periphery.

The regulatory mechanism of microtubule nucleation in tunicamycin-treated cells is unknown. However, C53 likely plays a role in this process since exogenous C53 decreased in the whole cell, including the centrosome in tunicamycin-treated cells. Moreover, endogenous C53 relocated from the subcellular fraction containing centrosomes after tunicamycin treatment. Thus the relocation of C53 from the centrosome could unblock microtubule nucleation in cells under ER stress. UFL1-catalyzed ufmylation is vital for relieving ER stress via ER-phagy (Liang e al., 2020). An increase in centrosomal microtubule nucleation might facilitate increased autophagic flux, ER expansion, and relief of ER stress.

In conclusion, in this study, we show that UFL1 and C53 interacting with γTuRC proteins play an important role in microtubule nucleation. C53, whose protein level is modulated by UFL1, associates with centrosome and represents a negative regulator of microtubule nucleation from the centrosomes. We demonstrate that the ER stress generated either by UFL1/C53 deletion or by pharmacological stressor stimulates centrosomal microtubule nucleation. The interaction of ER with newly formed microtubules could promote its enlargement to restore ER homeostasis. This suggests a novel mechanism for facilitating the ER network expansion under stress conditions.

## Material and methods

### Reagents

Nocodazole, puromycin, geneticin (G418), dimethyl pimelimidate dihydrochloride and all-*trans*-retinoic acid were from Sigma-Aldrich (St. Louis, MO). Lipofectamine 3000 was purchased from Invitrogen (Carlsbad, CA). Protein A Sepharose CL-4B and Glutathione Sepharose 4 Fast Flow were from GE Healthcare Life Sciences (Chicago, IL). Protease-inhibitor mixture tablets (Complete EDTA-free) were from Roche Molecular Biochemicals (Mannheim, Germany). Restriction enzymes were from New England Biolabs (Ipswich, MA). Oligonucleotides were synthesized by Sigma-Aldrich. Oligopeptides EYHAATRPDYISWGTQ (human γ-tubulin amino acid [aa] sequence 434-449) and EEFATEGTDRKDVFFY (human γ-tubulin aa sequence 38-53) (Zheng et al., 1991) were synthesized by Jerini Peptide Technologies (Berlin, Germany). ER-Tracker Green (BODIPY FL Glibenclamide) was from Molecular Probes (Eugene, OR), and 1mM stock was prepared in DMSO. Tunicamycin was from Sigma-Aldrich, and 1mg/ml stock was prepared in DMSO.

### Antibodie*s*

Catalog numbers for primary commercial Abs are shown in parentheses. Mouse mAbs TU-30 (IgG1) and TU-31 (IgG2b) to γ-tubulin aa sequence 434-449 were described previously (Nováková et al., 1996). Mouse mAbs GTU-88 to γ-tubulin aa sequence 38-53 (IgG1; T6557) and TUB2.1 to β-tubulin (IgG1, T4026), as well as rabbit Abs to actin (A2066), C53 (HPA022141), calnexin (C4731), DDRGK1 (HPA013373), GAPDH (G9545), GFP (G1544), histone H1.4 (H7665), PDI (P7496) and UFL1 (HPA030560; aa sequence 301-389) were from Sigma-Aldrich. Mouse mAbs to GCP4 (IgG1, sc-271876) and GCP6 (IgG1, sc-374063), as well as rabbit Abs to SHP-1 (sc-287) and p-Histone H3 (Ser 10) (sc-8656) were from Santa Cruz Biotechnology (Dallas, TX). Rabbit Abs to calcineurin (2614) and cyclin B1 (4138) were from Cell Signaling Technology (Danvers, MA). Mouse mAbs to C53 (IgG1; ab-57817) and DDIT3 (IgG2b, ab11419), as well as rabbit Ab to pericentrin (ab4448) were from Abcam (Cambridge, UK). Mouse mAb to GM130 (IgG1; 610822) was from BD Transduction Laboratories (San Jose, CA), rabbit Ab to GFP (11-476-C100) was from Exbio (Prague, Czech Republic), and rabbit Ab to pericentrin (ABT59) was from EMD-Millipore (La Jolla, CA). Rabbit Ab to α-tubulin (600-401-880) was from Rockland Immunochemicals (Limerick, PA).

Rabbit Ab to tRFP was from Evrogen (Moscow, Russia). Mouse mAbs GCP2-01 (IgG2b) and GCP2-02 (IgG1) to GCP2 were described previously (Dráberová et al., 2015), as well as mAb to α-tubulin TU-01 (Viklický et al., 1982). Rabbit Ab to non-muscle myosin heavy chain (BT-561; Biomedical Technologies., Stoughton, MA) and mAb NF-09 (IgG2a) to neurofilament NF-M protein (Dráberová et al., 1999) served as negative controls in the immunoprecipitation experiments. Rabbit Ab to GST was from Dr. Pe. Dráber (Institute of Molecular Genetics, CAS, Prague, Czech Republic).

Anti-mouse and anti-rabbit Abs conjugated with horseradish peroxidase (HRP) were from Promega Biotec (Madison, WI). TrueBlot anti-rabbit IgG HRP was purchased from Rockland Immunochemicals. Anti-mouse Abs conjugated with DyLight649, DyLight549, or AlexaFluor488 and anti-rabbit Abs conjugated with Cy3 or AlexaFluor488 were from Jackson Immunoresearch Laboratories (West Grove, PA).

Rabbit Ab to human UFL1 sequence 438-793 was prepared by immunizing three rabbits with purified GST-tagged human UFL1 fragment (GST-hUFL1_438-793). The procedure of immunization has been described previously (Dráber et al., 1991). Titers of sera were determined in ELISA using immobilized GST-hUFL1_438-793 or GST alone. Serum with the highest titer to GST-hUFL1_438-793 was partially purified and concentrated by ammonium sulfate precipitation. The precipitate was dissolved and dialyzed against PBS. Antigen or GST alone were covalently linked to Glutathione Sepharose 4 Fast Flow by dimethyl pimelimidate dihydrochloride as described (Bar-Peled and Raikhel, 1996). Prepared carriers were thereafter used for affinity isolation of Ab to UFL1. Shortly, a partially purified Ab was first preabsorbed with immobilized GST to remove anti-GST Abs. Then, Ab was bound onto the carrier with GST-hUFL_438-793. The solution of 0.1 M glycine-HCl, pH 2.5 was used for elution, and 0.2 ml fractions were immediately neutralized by adding 20 μl of 1M Tris-Cl, pH 8.0. Affinity-purified Ab stained GST-UFL1, but not GST alone, on the immunoblot, and detected UFL1 in whole-cell lysates.

### Cell cultures

Human osteogenic sarcoma cell line U-2 OS (U2OS) (Catalog No. ATCC, HTB-96), human glioblastoma cell line T98G (Catalog No. ATCC, CRL-1690), and human cervix adenocarcinoma HeLa S3 (Catalog. No. ATCC, CCL-2.2) were obtained from the American Type Culture Collection (Manassa, VA). Cells were grown at 37°C in 5% CO_2_ in DMEM supplemented with 10% FCS and antibiotics. P19.X1 cells, a subclone of mouse embryonal carcinoma cells P19, were cultured and subsequently differentiated by incubating cells with 1 μM all-*trans*-retinoic acid for nine days as described (Kukharskyy et al., 2004).

For ER live-cell imaging, cells were incubated with 1 μM ER-Tracker Green in Hank’s Balanced Salt Solution with calcium and magnesium (HBSS/Ca/Mg) for 30 min at 37°C. The staining solution was replaced with a probe-free medium and viewed by fluorescence microscopy. In some cases, cells were treated with 1 μg/ml tunicamycin or carrier (DMSO) for 24 h.

### DNA constructs

To prepare N-terminally EGFP-tagged human UFL1 *(UFL1;* Ref ID: NM_015323.4), the coding sequence was amplified by PCR from pF1KA0776 plasmid (Kasuza DNA Research Institute, Japan), containing the full-length cDNA of human UFL1. The following forward and reverse primers containing sites recognized by *KpnI/BamHI* restriction endonucleases (underlined) were used: forward 5’-AC<u>GGTACC</u>ATGGCGGACGCCT-3’ and reverse 5’-GGT<u>GGATCC</u>TTACTCTTCCGTCACAGATGA-3’. The PCR product was ligated to pEGFP-C1 vector (Clontech Laboratories, Mountain View, CA), resulting in plasmid pEGFP-hUFL1. To prepare C-terminally TagRFP-tagged human UFL1, the coding sequence without stop codon was amplified by PCR from pF1KA0776 plasmid. The following primers containing sites recognized by *NheI/SalI* were used: forward 5’-ATT<u>GCTAGC</u>AGAACCATGGCGGACGCCT-3’ and reverse 5’-CGC<u>GTCGA</u>CCTCTTCCGTCACAGATGATT-3’. The PCR product was ligated to pCI-TagRFP vector (Vinopal et al., 2012), resulting in plasmid phUFL1-TagRFP. To prepare N-terminally GST-tagged fulllength UFL1 (aa 1-794), the coding sequence was amplified from pF1KA0776 plasmid by PCR. The following primers containing sites recognized by *BamHI/SalI* were used: forward 5’-ATA<u>GGATCC</u>ATGGCGGACGCCTGG-3’and reverse 5’-CGAG<u>TCGAC</u>TTACTCTTCCGTCACAGATGATT-3’. The PCR product was ligated into pGEX-6P-1 (Amersham Biosciences, Uppsala, Sweden) using *BamHI/SalI* restriction sites, resulting in plasmid pGST-hUFL1_1-794. To generate Ab to human UFL1, the GST-tagged C-terminal fragment (aa 438-793) of UFL1 was prepared. The coding sequence was amplified from plasmid pF1KA0776 by PCR using the following primers containing sites recognized by *Bam*HI*/Sal*I: forward 5’-ATA<u>GGATCC</u>GGCAATGCCAGAGAG-3’; reverse 5’-TATC<u>GTCGA</u>CTCACTCTTCCGTCACAG-3. The PCR product was ligated into pGEX-6P-1 using *BamHI/SalI* restriction sites, resulting in plasmid pGST-hUFL1_438-793.

To prepare C-terminally EGFP-tagged human CDK5RAP3 *(CDK5RAP3;* Ref ID: NM_176096.3), the coding sequence without stop codon was amplified by PCR from the Myc-DDK-tagged CDK5RAP3 (tv3) plasmid (Origene Technologies, Rockville, MD; Catalog No. RC209901). The following primers containing sites recognized by *NheI/SalI* were used: forward 5’-T<u>GCTAGC</u>GGAGGAAAGATGGAGGAC-3’ and reverse 5’ T<u>GTCGAC</u>CAGAGAGGTTCCCATCAG-3’. The PCR product was ligated into pCR2.1 vector (Invitrogen) using *NheI/SalI* sites, resulting in plasmid pCR-hCDK5RAP3. The complete sequence of CDK5RAP3 without the stop codon was excised from pCR-hCDK5RAP3 by *NheI/Sal*I and ligated into pEGFP-N3 (Clontech), resulting in plasmid phCDK5RAP3-EGFP. To prepare C-terminally TagRFP-tagged human CDK5RAP3, the coding sequence without stop codon was excised from pCR-hCDK5RAP3 by *NheI/SalI* and inserted into pCI-TagRFP (Vinopal et al., 2012), resulting in plasmid phCDK5RAP3-TagRFP. Plasmid pGST-hCDK5RAP3 encoding N-terminally GST-tagged full-length human CDK5RAP3 was described previously (Hořejší et al., 2012).

To prepare mNeonGreen-tagged lentiviral vector with puromycin resistance, the coding sequence of mNeonGreen was digested out from the mNeonGreen-EB3-7 plasmid (Allele Biotechnology, San Diego, CA) by *Bam*HI*/Not*I. It was thereafter inserted into pCDH-CMV-MCS-EF1-puro vector (System Biosciences, Palo Alto, CA), resulting in vector mNeonGreen-puro. To prepare C-terminally mNeonGreen-tagged human γ-tubulin 1, the coding sequence was excised from pH3-16 plasmid (Zheng et al., 1991) by *NheI/EcoRI* and ligated into mNeonGreen-puro, resulting in vector phTUBG1-mNeonGreen-puro. Plasmid pGST-hTUBG1 encoding N-terminally GST-tagged full-length human γ-tubulin-1 was described previously (Macurek et al., 2008).

To prepare control C-terminally TagRFP-tagged mouse SHP-1, the coding sequence was excised from pmSHP-1-EGFP plasmid (Klebanovych, 2019) by *EcoRI/SalI* and ligated into pCI-Tag-RFP (Vinopal et al., 2012), resulting in plasmid pmSHP-1-TagRFP.

CRISPR/Cas9 gene editing (Sander and Joung, 2014) was used to disrupt the expression of human UFL1 (Ensembl: ENSG00000014123) or all human CDK5RAP3 variants (<u>Ensembl: ENSG00000108465</u>). Plasmids SpCas9 and pU6-sgRNAnew-III (donated by Dr. R. Malík, Institute of Molecular Genetics, CAS, Prague, Czech Republic) were used for optimal production of Cas9 and single-guide RNA (sgRNA), respectively. The CRISPR tool (https://zlab.bio/guide-design-resources) was used to design the DNA oligonucleotides (for production of sgRNA) that were cloned into *BsmBI* sites of pU6-sgRNAnew-III. To enrich for cells with disrupted expression of UFL1 or CDK5RAP3, we used the pRR-puro plasmid with a multiple cloning site that encodes a non-functional puromycin resistance cassette (Flemr and Buhler, 2015). Annealed sense and anti-sense oligonucleotides containing the sequences from the region of interest and overhangs with *AatII/SacI* restriction sites were ligated into pRR-puro digested with *AatII/SacI,* resulting in reporter plasmids pRR-hUFL1-puro or pRR-hCDK5RAP3-puro. Co-transfection of the reporter plasmid with the plasmids encoding sgRNAs and Cas9 led to CRISPR-induced double-strand break (DSB) in the reporter plasmid. When the DSB was repaired by homologous recombination, the puromycin resistance was restored.

### Generation of UFL-1 and CDK5RAP3 deficient cell lines

In order to delete part of the 5’ region of the UFL1 gene, U2OS cells were transfected with CRISPR/Cas9 vectors (sgRNA#1, sgRNA#2, SpCas9) together with reporter plasmid pRR-hUFL1-puro by transfection using Lipofectamine 3000 according to the manufacturer’s instructions. The final transfection mixture in a 24-well plate contained 200 ng of sgRNA#1, 200 ng of sgRNA#2, 200 ng of reporter plasmid, and 400 ng of SpCas9 in 1 ml of Dulbecco’s modified Eagle’s medium (DMEM) containing 10% FCS, penicillin (100 units/ml), and streptomycin (0.1 mg/ml). The medium was changed after 48 h, and puromycin was added to the final concentration of 2 μg/ml. The stable selection was achieved by culturing cells for 1 wk in the presence of puromycin. The single-cell dilution protocol (Green and Sambrook, 2012) was used to obtain cell clones that were thereafter analyzed by PCR and immunoblotting. Single-cell clones were expanded, genomic DNA was extracted with the QIAamp DNA Mini Kit (QIAGEN, Gilden, Germany), and deletion in the UFL1 gene was determined by PCR amplification with primers flanking the deleted region: forward 5’-AGGCGCCAATCTTAGACACAG-3’; reverse 5’-CAAAAGCTGCCCTTTTATCTGT-3’. Amplified fragments were visualized in 2% agarose gels stained by GelRed Nucleic Acid Gel Stain (Biotium, Fremont, CA). While amplification of short fragments (~ 570 bp) was detected in UFL1 deficient clones, no amplification was found in control U2OS due to the large size of the deleted region (~ 7 kb). The PCR fragments were subcloned into pCR2.1 vector (Invitrogen), and individual colonies were sequenced.

A similar approach was applied to the preparation of cells deficient in CDK5RAP3. In order to delete part of the 5’ region of the GDK5RAP3 gene containing the canonical and alternative start codons, U2OS cells were transfected with CRISPR/Cas9 vectors (sgRNA#1, sgRNA#2, SpCas9) together with reporter plasmid pRR-hCDK5RAP3-puro. Deletion in the CDK5RAP3 gene was determined by PCR amplification with primers flanking the deleted region: forward 5’-CATGCATCCATCATCCCAG-3’; reverse 5’-TGACATGTGACGTGTGAAACTCT-3’. Amplified fragments were visualized in agarose gels. While amplification of short fragments (~ 700 bp) was detected in CDK5RAP3 deficient clones, no amplification was found in control U2OS due to the large size of the deleted region (~ 6 kb). The PCR fragments were subcloned into pCR2.1 vector (Invitrogen), and individual colonies were sequenced.

### Generation of cell lines expressing tagged proteins

To prepare U2OS cells expressing EGFP-, TagRFP-or mNeonGreen-tagged proteins, cells were transfected with 2.5 μg DNA per 3-cm tissue culture dish using Lipofectamine 3000 according to the manufacturer’s instructions. After 12 h, the transfection mixture was replaced with a complete fresh medium, and cells were incubated for 48 h. Cells were thereafter incubated for one wk in complete fresh medium containing G418 at concentration 1.2 mg/ml. In the case of phTUBG1-mNeonGreen-puro transfection, cells were incubated in the presence of puromycin at a concentration of 2 μg/ml. For phenotypic rescue experiments, cells expressing TagRFP-tagged proteins were flow-sorted using a BD Influx cell sorter (BD Bioscience, San Jose, CA). TagRFP emission was triggered by 561 nm laser; fluorescence was detected with a 585/29 bandpass filter.

### Real-time qRT-PCR

Total RNA from control or tunicamycin-treated cells was isolated with the RNeasy Mini Kit (QIAGEN, Gilden, Germany) and converted to cDNA using the High-Capacity cDNA Reverse Transcription Kit (Applied Biosystems, Waltham, MA) according to the manufacturer’s protocol. The quantitative PCRs were performed with primers specific for human calnexin *(CANX;* forward 5’-GAACATTCTTCCCTTTGACC-3’ and reverse 5’-TCTTAGAGTTCTCATCTGGAC-3’; primers anneal to all transcript variants), DNA damage-inducible transcript 3 *(DDIT3;* forward 5’-AACATCACTGGTGTGGAGGC-3’ and reverse 5’-TGCACAGTTCAGCGGGTA-3’; primers anneal to all transcript variants except NM_001195057.1), and prolyl 4-hydroxylase subunit beta (protein disulfide-isomerase; *P4HB;* forward 5’-CTTAAGGAGCTTATTGAGGAG-3’ and reverse 5’-CATTGCAATTGGAGATGTTG-3’). Peptidylprolyl isomerase A (cyclophilin A; *PPIA;* forward 5’-GAGCACTGGGGAGAAAGGAT-3’ and reverse 5’-CTTGCCATCCAGCCACTCAG-3’; primers anneal to all transcript variants) served as an internal control. All primers were tested in silico by the Basic Local Alignment Search Tool from the National Center for Biotechnology Information (BLAST NCBI; NIH, Bethesda, MD) to amplify the specific targets. Quantitative PCRs were performed in a LightCycler 480System (Roche, Mannheim, Germany) as described (Dráberová et al., 2017). Each sample was run in triplicate. The identity of the PCR products was verified by sequencing.

### Preparation of cell extracts

Whole-cell lysates for SDS-PAGE were prepared by washing the cells in cold HEPES buffer (50 mM HEPES pH 7.6, 75 mM NaCl, 1 mM MgCl_2_, and 1 mM EGTA), solubilizing them in hot SDS-sample buffer, and boiling for 5 min. When preparing whole-cell extracts for immunoprecipitation and GST pull-down assays, cells grown on a 9-cm Petri dish (cell confluence 80%) were rinsed in HEPES buffer and extracted (0.7 ml/Petri dish) for 10 min at 4°C with HEPES buffer supplemented with protease and phosphatase (1 mM Na3VO4 and 1 mM NaF) inhibitors and 1% NP-40 (extraction buffer). The suspension was then spun down (12,000 x g, 15 min, 4°C), and the supernatant was collected. When preparing whole-cell extracts for gel filtration chromatography, cells grown on six 9-cm Petri dishes were scraped into cold HEPES buffer, and pelleted cells were extracted in 1 ml of extraction buffer.

To prepare the crude membranous fraction, cells from five 9-cm Petri dishes were released by trypsin, washed in HEPES, and mechanically disrupted in 3.5 ml of cold HEPES buffer supplemented with inhibitors using a Dounce homogenizer (disruption efficiency was verified under a microscope). The homogenate was centrifuged at 300 x g for 5 min (supernatant S1, pellet P1). Post-nuclear supernatant was centrifuged at 1,400 x g for 10 min (supernatant S2, pellet P2). Pelleted material (P2; crude membranous fraction) was solubilized for 5 min at 4°C with 1.4 ml of extraction buffer, and the suspension was spun down (12,000 x g, 15 min, 4°C). The supernatant was collected for immunoprecipitation experiments. For gel filtration chromatography, membranes prepared from twelve 9-cm Petri dishes were solubilized in 1 ml of extraction buffer.

### Gel filtration chromatography

Gel filtration was performed using fast protein liquid chromatography (AKTA-FPLC system) on a Superose 6 10/300 GL column (GE Healthcare Life Sciences) as described previously (Hořejší et al., 2012). The column equilibration and chromatography were performed in HEPES buffer, and 0.5-ml aliquots were collected. Samples for SDS-PAGE were prepared by mixing with 5x concentrated SDS-sample buffer.

### Immunoprecipitation, GST pull-down assay, gel electrophoresis, and immunoblotting

Immunoprecipitation was performed as previously described (Blume et al., 2008). Extracts were incubated with beads of protein A saturated with mouse mAbs to (i) γ-tubulin (TU-31; IgG2b), (ii) GCP2 (GCP2-01; IgG2b), (iii) NF-M (NF-09; IgG2a, negative control), or with rabbit Abs to (iv) UFL1_301-389_, (v) UFL1_438-793_, (vi) C53 (Sigma-Aldrich), (vii) (GFP), (viii) non-muscle myosin (negative control) or with (ix) immobilized protein A alone. Antibodies C53, GFP, UFL1_301-389_, and UFL1_438-793_, were used at Ig concentrations 0.5-2.5 μg/ml. Ab to myosin was used at a dilution of 1:100. The mAbs TU-31, GCP2-01, and NF-09, in the form of hybridoma supernatants, were diluted 1:2.

To identify proteins interacting with membrane-bound γ-tubulin, 1% NP-40 extract from the microsomal fraction (P100) of differentiated P19 cells (Macurek et al., 2008) was incubated with anti-peptide mAb TU-31 immobilized on protein A carrier. After extensive washing, the bound proteins were eluted with immunizing peptide EYHAATRPDYISWGTQ (human γ-tubulin, aa sequence 434-449) (Nováková et al., 1996) as described (Hořejší et al., 2012). Peptide EEFATEGTDRKDVFFY (human γ-tubulin, aa sequence 38-53) served as a negative control.

Preparation and purification of GST-tagged fusion proteins were described previously (Frangioni and Neel, 1993), as were pull-down assays with whole-cell extracts (Kukharskyy et al., 2004).

Gel electrophoresis and immunoblotting were performed using standard protocols (Caracciolo et al., 2010). For immunoblotting, mouse mAbs to γ-tubulin (GTU-88), GCP4, GCP6, DDIT3 and GM130 were diluted 1:10,000, 1:1,000, 1:500, 1:500 and 1:250, respectively. Mouse mAbs to α-tubulin (TU-01) and GCP2 (GCP2-02), in the form of spent culture supernatant, were diluted 1:10 and 1:5, respectively. Rabbit Abs to actin, GFP (Sigma-Aldrich), UFL1 (Sigma-Aldrich), C53 (Sigma-Aldrich), tRFP, cyclin B1, calcineurin, and p-Histone H3 were diluted 1:10,000, 1:5,000, 1:3,000, 1:3,000, 1:2,000, 1:1,000, 1:1,000, and 1:700, respectively. Rabbit Abs to PDI, histone H1.4, pericentrin (EMD-Millipore) and DDRGK1 were diluted 1:50,000, 1:1,000, 1:1,000 and 1:500, respectively. Rabbit Abs to calnexin, GAPDH, GST, and SHP-1 were diluted 1:100,000. Secondary anti-mouse and anti-rabbit Abs conjugated with HRP were diluted 1:10,000. TrueBlot anti-rabbit IgG HRP was diluted 1:100,000. The HRP signal was detected with SuperSignal WestPico or Supersignal West Femto Chemiluminescent reagents from Pierce (Rockford, IL) and the LAS 3000 imaging system (Fujifilm, Düsseldorf, Germany). The AIDA image analyzer v5 software (Raytest, Straubenhardt, Germany) was used to quantify signals from the immunoblots.

### Mass spectrometry

Concentrated samples after peptide elution were dissolved in 2x Laemmli sample buffer and separated in 8% SDS-PAGE. Gels were stained with Coomassie Brilliant Blue G-250. The bands of interest were excised from the gel, destained, and digested by trypsin. The extracted peptides were analyzed by a MALDI-TOF mass spectrometer (Ultraflex III; Bruker Daltonics, Bremen, Germany) in a mass range of 700-4000 Da and calibrated internally using the monoisotopic [M+H]^+^ ions of trypsin auto-proteolytic fragments (842.51 and 2211.10 Da). Data were processed using FlexAnalysis 3.3 software (Bruker Daltonics, Bremen Germany) and searched by the in-house Mascot search engine against the SwissProt database subset of all *Mus musculus* proteins.

### Evaluation of cell growth

Cell proliferation was assessed by manual cell counting of control U2OS, UFL1_KO or C53_KO cells. A total of 2 x 10^5^ cells diluted in culture medium were plated on a 6-cm-diameter Petri dish. Cells were counted at various time intervals from 1 to 4 days. Samples were counted in doublets in a total of three independent experiments.

### Microtubule regrowth experiments

Microtubule regrowth from centrosomes was followed in a nocodazole washout experiment. Cells growing on coverslips were treated with nocodazole at a final concentration of 10 μM for 90 min at 37 °C to depolymerize microtubules. Cells were then washed with phosphate-buffered saline (PBS) precooled to 4 °C (3 times 5 min each) to remove the drug, transferred to complete medium tempered to 28 °C, and microtubule regrowth was allowed for 1-3 min at 28 °C. Cells were after that fixed in formaldehyde and extracted in Triton X-100 (F/Tx) and postfixed in cold methanol (F/Tx/M).

### Immunofluorescence microscopy

Cells were fixed and immunostained as described (Dráberová and Dráber, 1993). Samples were fixed in F/Tx, and for double-label experiments with anti-γ-tubulin Ab, they were postfixed in methanol (F/Tx/M). To visualize TagRFP-tagged proteins, cells were permeabilized with 10 μM digitonin in CHO buffer (25 mM HEPES, 2 mM EGTA, 115 mM CH_3_COOK, 2.5 mM MgCl_2_, 150 mM sucrose, pH 7.4) for 30 sec, then fixed with 3% formaldehyde in microtubule-stabilizing buffer (MSB) (Dráberová and Dráber, 1993) for 20 min at room temperature and postfixed in methanol at −20°C for 5 min (D/F/M). Alternatively, TagRFP-tagged proteins were visualized in samples extracted in 0.5% Triton X-100 for 1 min at room temperature, then fixed with 3% formaldehyde in MSB for 20 min at room temperature, and postfixed in methanol (Tx/F/M). Mouse mAbs to β-tubulin (TUB2.1), C53 (Abcam), and DDIT3 were diluted 1:500, 1:200, and 1:50, respectively. Rabbit Abs to calnexin, PDI, pericentrin, and UFL1_438-793_ were diluted 1:1,000, 1:1,000, 1:250, and 1:50, respectively. Mouse mAb to γ-tubulin (TU-30), in the form of spent culture supernatant, and rabbit Ab to α-tubulin were diluted 1:10 and 1:100, respectively. Secondary AlexaFluor488-, DyLight549- and DyLight649-conjugated anti-mouse Abs were diluted 1:200, 1:1,000 and 1:1,000, respectively. The AlexaFluor488- and Cy3-conjugated anti-rabbit Abs were diluted 1:200 and 1:1,000, respectively. Samples were mounted in MOWIOL 4-88 (Calbiochem, San Diego, CA) and examined with an Olympus AX-70 Provis microscope (Olympus, Hamburg Germany) equipped with a 60×/1.0 NA water objective or with a Delta Vision Core system (AppliedPrecision, Issaquah, WA) equipped with a 60×/1.42 NA oil objective.

To quantify the microtubule regrowth, different areas per sample were taken in both fluorescence channels. The sum of γ-tubulin or α-tubulin immunofluorescence intensities was obtained from nine consecutive frames (0.2 μm steps), with the middle frame chosen with respect to the highest γ-tubulin intensity. Quantification of the microtubule regrowth assay was analyzed automatically in 2-μm regions of interest (ROIs) centered at the centrosomes, marked by γ-tubulin staining, using an in-house written macro for Fiji processing program (Klebanovych et al., 2019; Schindelin et al., 2012).

### Microtubule nucleation visualized by time-lapse imaging

For time-lapse imaging, U2OS cells expressing EB3-mNeonGreen were grown on a 35 mm μ-Dish with ibidi polymer coverslip bottom (Ibidi GmbH, Gräfelfing, Germany). Prior to imaging, the medium was replaced for FluoroBrite™ DMEM (ThermoFisher Scientific, Waltham, MA), supplemented with 25 mM HEPES and 1% FCS, 30 min before imaging. Time-lapse sequences were collected in seven optical slices (0.1 μm steps) for 1 min at 1 s interval with the Andor Dragonfly 503 spinning disc confocal system (Oxford Instruments, Abingdon, UK) equipped with a stage top microscopy incubator (Okolab, Ottaviano, Italy), 488 nm solid-state 150 mW laser, HCX PL APO 63x oil objective, NA 1.4, and Zyla sCMOS 16 bit camera. For each experiment, at least 10 cells were imaged (acquisition parameters: 40 μm pinhole size, 15% laser power, 50 ms exposure time, 525/50 nm emission filter). The time-lapse sequences were deconvoluted with Huygens Professional software v. 19.04 (Scientific Volume Imaging, Hilversum, the Netherlands), and maximum intensity projection of z stack was made for each time point in Fiji. Newly nucleated microtubules were detected by manual counting of EB3 comets emanating from the centrosomes.

### Statistical analysis

A minimum of three independent experiments was analyzed for each quantification. The counts of individual data points were indicated in the figure legends. All data were presented as mean ± SD. Significance was tested using a two-tailed, unpaired Student’s *t* test or one-way ANOVA followed by a Sidak’s post hoc test using Prism 8 software (GraphPad Software, San Diego). The used test are indicated in the figure legends. For all analyses,*p* values were represented as follows: *,*p* < 0.05; **,*p* < 0.01; ***,*p* < 0.001; ****,*p* < 0.0001

### On line supplemental material

Table S1 shows mass spectrometry identification of UFL1. Fig S1 shows results from the additional experiments supplementing Fig. 1, and controls for immunoprecipitation experiments. Fig. S2 documents preparation and properties of *UFL1 and CDK5RAP3* knockout cell lines. Fig. S3 shows that deletion of UFL1 and C53 stimulates accumulation of γ-tubulin but not pericentrin at the centrosome, and promotes microtubule nucleation. Fig. S4 presents the distribution of PDI in knockout cell lines, expression of calnexin in phenotypic rescue experiment and upregulation of ER stress-associated proteins after tunicamycin treatment. List of source data summarizes figure panels for which source data are provided (in Fig. 4, Fig. 5, Fig. 6, Fig. 7, Fig. 8, Fig. 9, Fig. S3 and Fig. S4).

## Acknowledgements

We thank Dr. B. R. Oakley (University of Kansas, Lawrence, KS, USA) for the pH3-16 vector encoding human TUBG1; Drs. R. Malík and P. Svoboda (Institute of Molecular Genetics, CAS, Prague, Czech Republic) for providing plasmids spCas9, pU6-sgRNAnew-III and pRR-puro; and Dr. Pe. Dráber (Institute of Molecular Genetics, CAS, Prague, Czech Republic) for providing polyclonal Ab to GST. We also thank I. Mlchová for the excellent technical assistance.

## Abbreviations

C53: CDK5RAP3
DDIT3: DNA damage-inducible transcript 3 protein
DDRGK1: DDRGK domain-containing protein 1
GCP: γ-tubulin complex protein
MTOC: microtubule-organizing center
PDI: protein disulfide-isomerase
γTuRC: γ-Tubulin ring complex
UFL1: E3 UFM1-protein ligase 1
UFM: ubiquitin-fold modifier 1
UPR: unfolded protein response

## Supplemental material

**Table S1.** Mass spectrometry identification of E3 UFM1-protein ligase 1 (UFL1).

| Measured mass [M+H] | Computed mass [M+H] | Error (ppm) | Peptide sequence | Peptide position |
| --- | --- | --- | --- | --- |
| 1292.619 | 1292.605 | 11 | MoxADAWEEIRR | 1 - 10 |
| 820.454 | 820.431 | 28 | LAADFQR | 11 - 17 |
| 1037.488 | 1037.501 | 13 | AQFAESTQR | 18 - 26 |
| 2369.293 | 2369.271 | 9 | QLEVVHTLDGKEYITPAQISK | 44 - 64 |
| 2305.286 | 2305.251 | 15 | VNIVDLQQVINVDLTHIESR | 77 - 96 |
| 1808.942 | 1808.918 | 13 | AYDLPGDFLTQALTQR | 145 - 160 |
| 1279.699 | 1279.675 | 19 | IINGHLDLNR | 164 - 174 |
| 1209.630 | 1209.663 | 27 | GVIFTEAFVAR | 175 - 185 |
| 2317.226 | 2317.171 | 24 | YGFQEQLLYSVLEDLVSTGR | 210 - 229 |
| 1166.597 | 1166.620 | 20 | AVFVPDIYSR | 242 - 251 |
| 1373.633 | 1373.648 | 11 | TQSTWVDSFFR | 252 - 262 |
| 1412.713 | 1412.681 | 23 | QNGYLEFDALSR | 263 - 274 |
| 1445.695 | 1445.673 | 15 | FITDCcamTGLFSER | 368 - 379 |
| 1521.752 | 1521.791 | 26 | NNPVHLITEEDLK | 390 - 402 |
| 2049.112 | 2049.061 | 25 | TIKDLQEEVSNLYNNIR | 535 - 551 |
| 1706.800 | 1706.834 | 20 | DLQEEVSNLYNNIR | 538 - 551 |
| 941.514 | 941.532 | 19 | QILFQHR | 662 - 668 |

Mox: oxidized methionine; Ccam: carbamidomethylated cysteine
1 **MADAWEEIRR** **LAADFQRAQF** **AESTQRL**SER NCIEIVNKLI SQ**KQLEVVHT** **LDGKEYITPA** 61 **QISKEMRDEL** HVRGGR**VNIV** **DLQQVINVDL** **THIESRVSDI** IKSEKHVQMV LGQLIDENYL 121 DQLSEEVNDK LQESGQVTVS ELCK**AYDLPG** **DFLTQALTQR** LGR**IINGHLD** **LDNRGVIFTE** 181 **AFVAR**HKARI RGLFSAITRP TPNVNSLVSKY **GFQEQLLYSV** **LEDLVSTGRL** RGTVVGGRQD 241 **KAVFVPDIYS** **RTQSTWVDSF** **FRQNGYLEFD** **ALSRL**GIPDA VNYIKKRYKN TQLLFLKATC 301 VGQGLVDQVE ASVEEAISSG TWVDISPLLP SSLSVEDAAM LLQQVMRPFQ KLASAIVFSD 361 TVVVSEK**FIT** **DCTGLFSER**M HQKAEKEMKN **NPVHLITEED** **LKQ**ISILESV NTSKKDKKDE 421 RRKKATEGSG SVRGGGGGNA REYKIKKTKK KGRKDEDSDD ESQSSHGGKK KPDITFMFQD 481 EIEDCLRKHI QDAPEEFISE LAEYLIKPLN KMYLEVVRVS FMSSTSASGT GRKR**TIKDLQ** 541 **EEVSNLYNNI** **RLF**EKGMKYF ADDTQTALTK HLLKTVCTDI TNLMFNFLAS DFLMAVEEPA 601 AITSDIRKKI LSKLTEETKV ALTKLHNSLN EKSIEDFLSC LDSATEACDI MVKKGDKKRE 661 **RQILFQHR**QA LCEQLKVTEd PALILHLTAV LLFQLSTHSM LHAPGRCVPQ IIAFLHSKIP 721 EDQHTLLVKY QGLVVKQLVS QNKKTGQGED PSSDELDKEQ HDVTNATRKE LQELSLSIKD 781 LVLKSRKSSV TEE
Amino acids of the identified peptides in the mouse E3 UFM1-protein ligase 1 (Uniprot identifier Q8CCJ3) are indicated in bold and underlined (bottom part of the table). The matched peptides cover 26% of the protein sequence.

**Figure S1.**
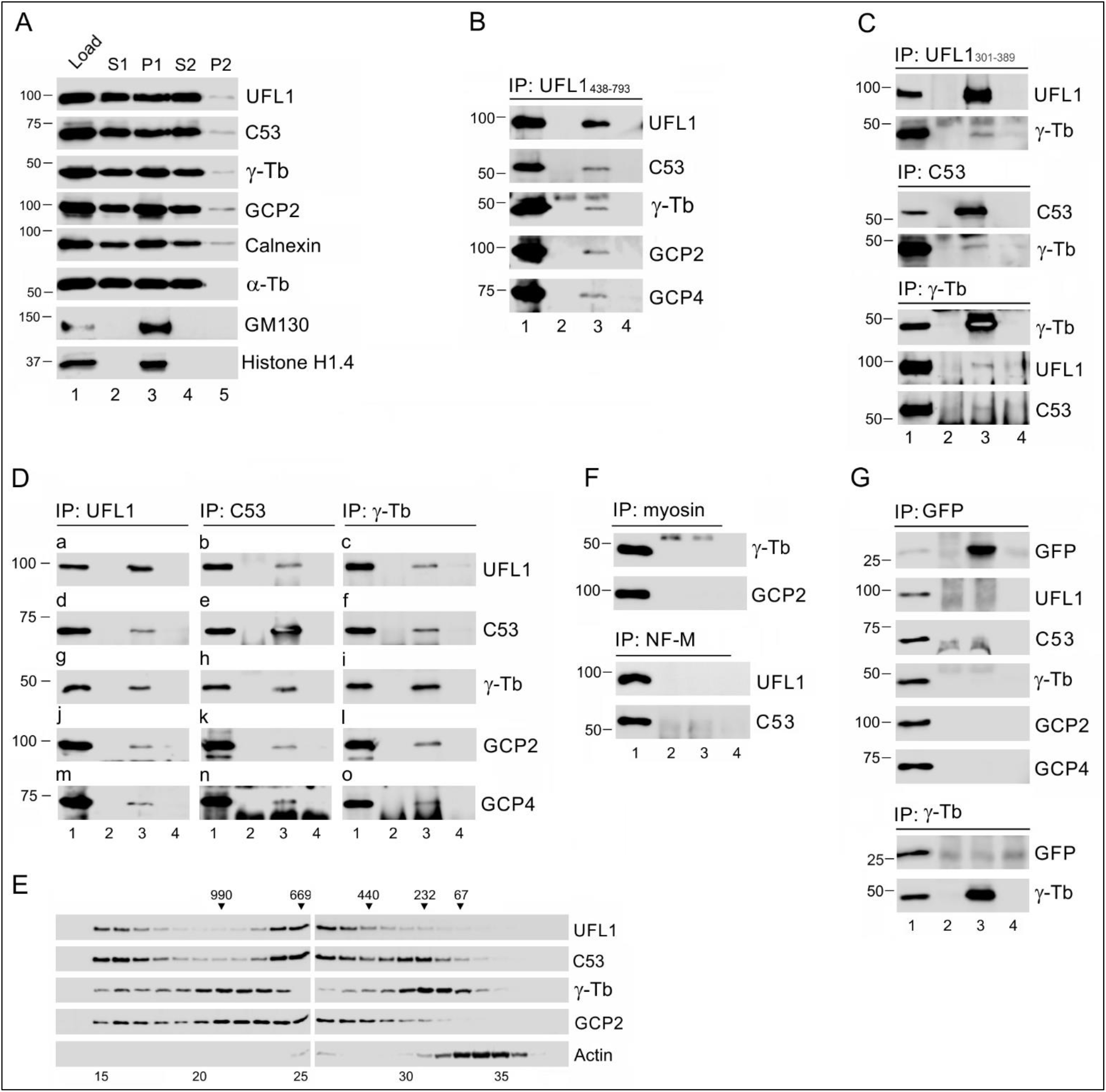
UFL1 and C53 associate with γTuRC proteins. Controls for immunoprecipitation experiments. (**A-C**) Membrane-bound UFL1 and C53 interact with γTuRC proteins. (**A**) Relative distribution of proteins in fractions after differential centrifugation of the U2OS cell homogenate. Cell fractions were prepared as described in the Materials and Methods section. Cell homogenate *(lane 1),* supernatant S1 *(lane 2),* pellet P1 *(lane 3),* supernatant S2 *(lane 4),* pellet P2 *(lane 5*). To compare the relative distribution of proteins, pelleted material was resuspended in a volume equal to that of the corresponding supernatant. Blots were probed with Abs to UFL1, C53, γ-tubulin (γ-Tb), GCP2, calnexin, α-tubulin (α-Tb), GM130 and Histone H1.4. (**B-C**) Immunoprecipitation experiments. Extracts from the membranous fraction (P2) of U2OS (**B**) or T98G (**C**) cells were precipitated with immobilized Abs specific to UFL1_438-793_ (**B**) or UFL1_301-389_, C53 and γ-tubulin (**C**). The blots were probed with Abs to UFL1, C53, γ-tubulin (γ-Tb), GCP2, and GCP4. Load *(lane 1*), immobilized Abs not incubated with cell extracts *(lane 2),* precipitated proteins *(lane 3*), and carriers without Abs and incubated with cell extracts *(lane 4).* (**D-E**). UFL1 and C53 in the whole-cell extract interacts with γTuRC proteins. (**D**) Immunoprecipitation experiments. Precipitation of U2OS whole-cell extracts with immobilized Abs specific to UFL1 (**a, d, g, j, m**), C53 (**b, e, h, k, n**), or γ-tubulin (**c, f, i, l, o**). The blots were probed with Abs to UFL1, C53, γ-tubulin (γ-Tb), GCP2, or GCP4. Load *(lane 1),* immobilized Abs not incubated with cell extracts *(lane 2),* precipitated proteins *(lane 3),* and carriers without Abs and incubated with cell extracts *(lane 4).* (**E**) The size distribution of proteins in U2OS whole-cell extracts fractionated on the Superose 6 column. The blots of the collected fractions were probed with Abs to UFL1, C53, γ-tubulin (γ-Tb), GCP2, and actin. The calibration standards (in kDa) are indicated on the top. The numbers at the bottom denote individual fractions. (**F-G**) Negative controls for immunoprecipitation experiments. (**F**) Isotype controls. Whole-cell extracts from the U2OS membranes were precipitated with immobilized rabbit Ab to myosin or mouse mAb to NF-M (IgG2a). Blots were probed with Abs to γ-tubulin (γ-Tb), GCP2, UFL1, or C53. Load *(lane 1*), immobilized Abs not incubated with cell extracts *(lane 2),* precipitated proteins (*lane 3*), and carriers without Abs and incubated with cell extracts (*lane 4*). (**G**) Whole-cell extracts from the U2OS cells expressing EGFP alone precipitated with immobilized Abs to GFP and γ-tubulin. The blots were probed with Abs to GFP, UFL1, C53, γ-tubulin (γ-Tb), GCP2, or GCP4. Load *(lane 1),* immobilized Abs not incubated with cell extracts (*lane 2*), precipitated proteins (*lane 3*), and carriers without Abs and incubated with cell extracts (*lane 4*).

**Figure S2.**
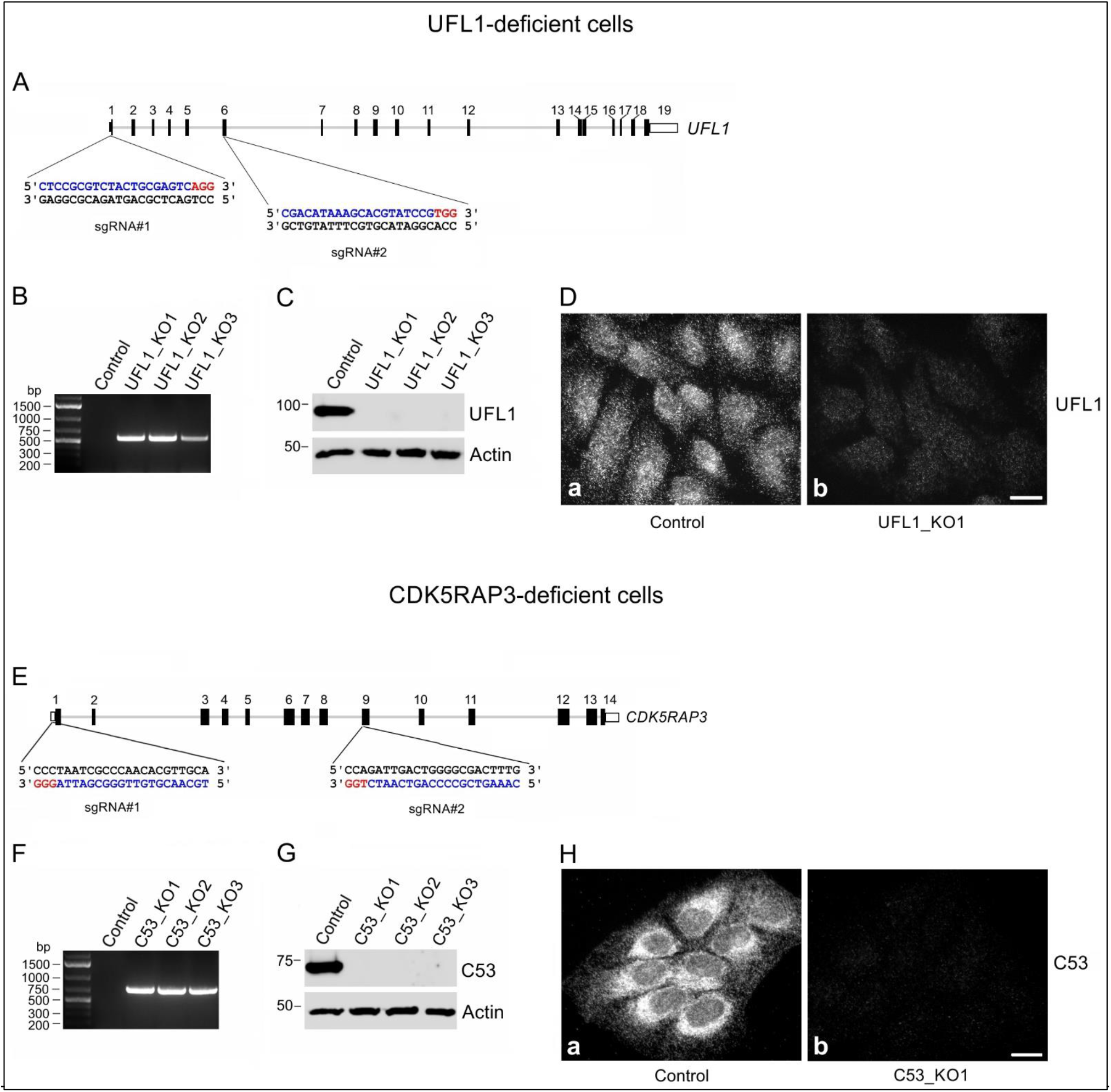
Generation of *UFL1* and *CDK5RAP3* knockout cell lines. (**A-D**) UFL1-deficient cells. (**A**) Schematic diagram of the longest transcript of *UFL1* gene (34.9 kb), containing 19 exons, with sites targeted by guide RNA (sgRNA) sequences. Targeted sites (blue) and protospacer adjacent motifs (PAM; red) are depicted. (**B**) PCR amplification of genomic DNA from control U2OS cells (Control) and UFL1-deficient U2OS cell lines (UFL1_KO1, UFL1_KO2, UFL2_KO3) with primers flanking the deleted region. Due to the large size of the deleted region (~6.8kb), no amplification was found in control cells. Amplification of short fragments (~560bp) was detected in UFL1-deficient clones. **(C)** UFL1 protein levels in U2OS and UFL1-deficient U2OS cell lines analyzed by immunoblotting of whole-cell lysates. Actin served as the loading control. (**D**) UFL1 protein levels in control (a) and UFL1_KO1 (**b**) cells analyzed by immunofluorescence microscopy with Ab to UFL1_438-793_. Fixation F/Tx. The pairs of images were collected and processed in the same manner. Scale bar, 20 μm. (**E-H**) CDK5RAP3-deficient cells. (**E**) Schematic diagram of the longest transcript of *CDK5RAP3* gene (15.2 kb), containing 14 exons, with sites targeted by guide RNA (sgRNA) sequences. Targeted sites (blue) and protospacer adjacent motifs (PAM; red) are depicted. (**F**) PCR amplification of genomic DNA from control U2OS cells (Control) and CDK5RAP3 (C53)-deficient U2OS cell lines (C53_KO1, C53_KO2, C53_KO3) with primers flanking the deleted region. Due to the large size of the deleted region (~6.1 kb), no amplification was found in control cells. Amplification of short fragments (~720bp) was detected in C53-deficient clones. **(G)** C53 protein levels in U2OS and C53-deficient U2OS cell lines analyzed by immunoblotting of whole-cell lysates. Actin served as the loading control. (**H**) C53 protein levels in control (**a**) and C53_KO1 (**b**) cells analyzed by immunofluorescence microscopy with Ab to C53. Fixation F/Tx. The pairs of images were collected and processed in the same manner. Scale bar, 20 μm.

**Figure S3.**
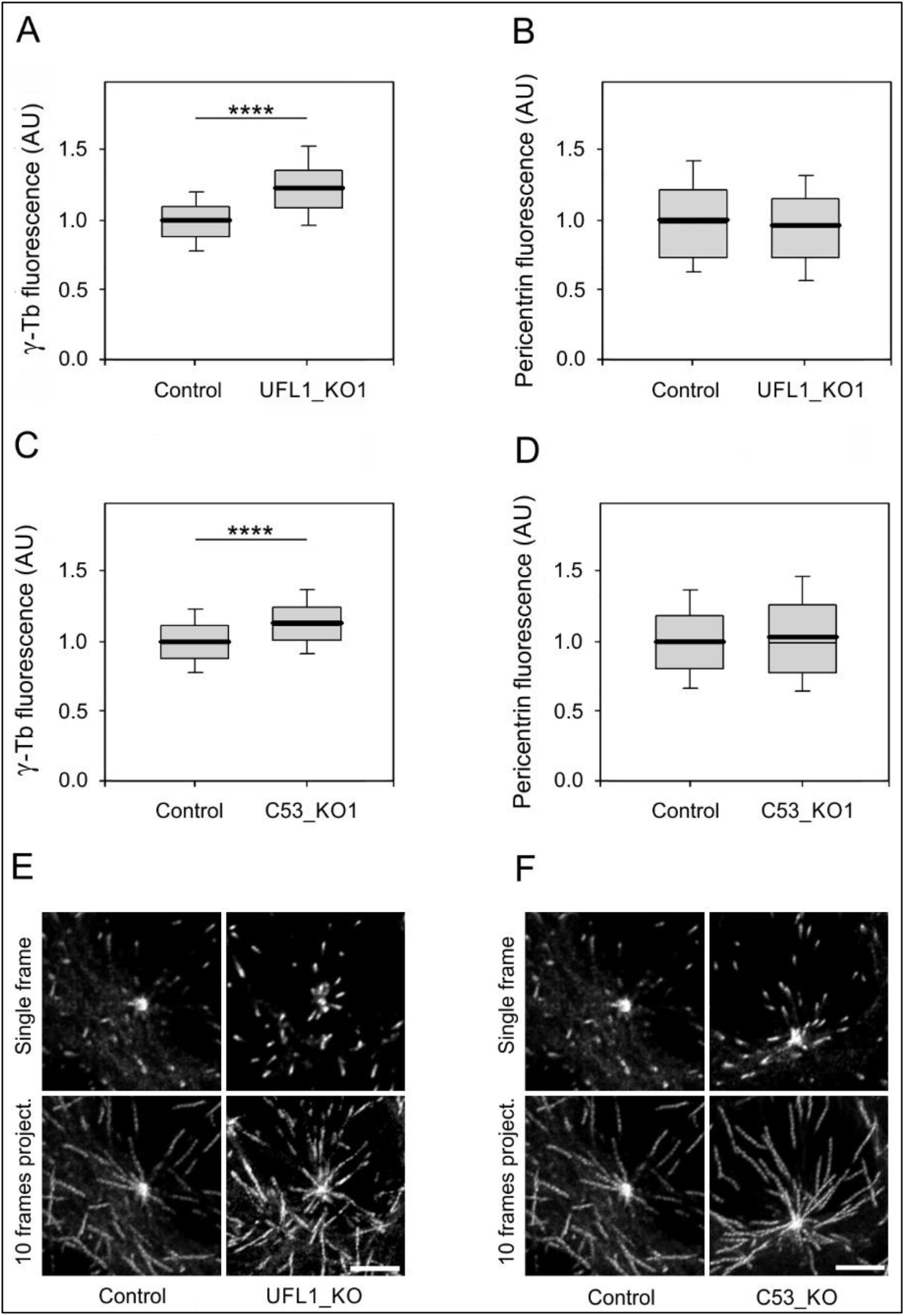
Deletion of UFL1 or C53 stimulates accumulation of γ-tubulin but not pericentrin at the centrosome, and promotes microtubule nucleation. (**A-D**) The distributions of γ-tubulin or pericentrin fluorescence intensities (arbitrary units [AU]) in 2-μm ROI at 2.0 min of microtubule regrowth in control, UFL1-deficient (UFL1_KO1) or C53-deficient (C53_KO1) cells are shown as box plots (three experiments for UFL1-KO1 and four experiments for C53_KO1, > 49 cells counted for each experimental condition). (**A-B**) Box plot of γ-tubulin (**A**) and pericentrin (**B**) fluorescence intensities in UFL1_KO1 cells (n=234) relative to control cells (Control, n=247). (**C-D**) Box plot of γ-tubulin (**C**) and pericentrin (**D**) fluorescence intensities in C53_KO1 cells (n=358) relative to control cells (Control, n=322). The bold and thin lines within the box represent mean and median (the 50th percentile), respectively. The bottom and top of the box represent the 25th and 75th percentiles. Whiskers below and above the box indicate the 10th and 90th percentiles. Two-tailed, unpaired Student’s *t* test was performed to determine statistical significance. ****, p < 0.0001. (**E-F**) Time-lapse imaging of control and UFL1_KO1 (**E**) or C53_KO1 (**F**) cells expressing EB3-mNeonGreen. Still images of EB3 (Single frame) and tracks of EB3 comets over 10 s (10 frames project.). Scale bar s, 5 μm.

**Figure S4.**
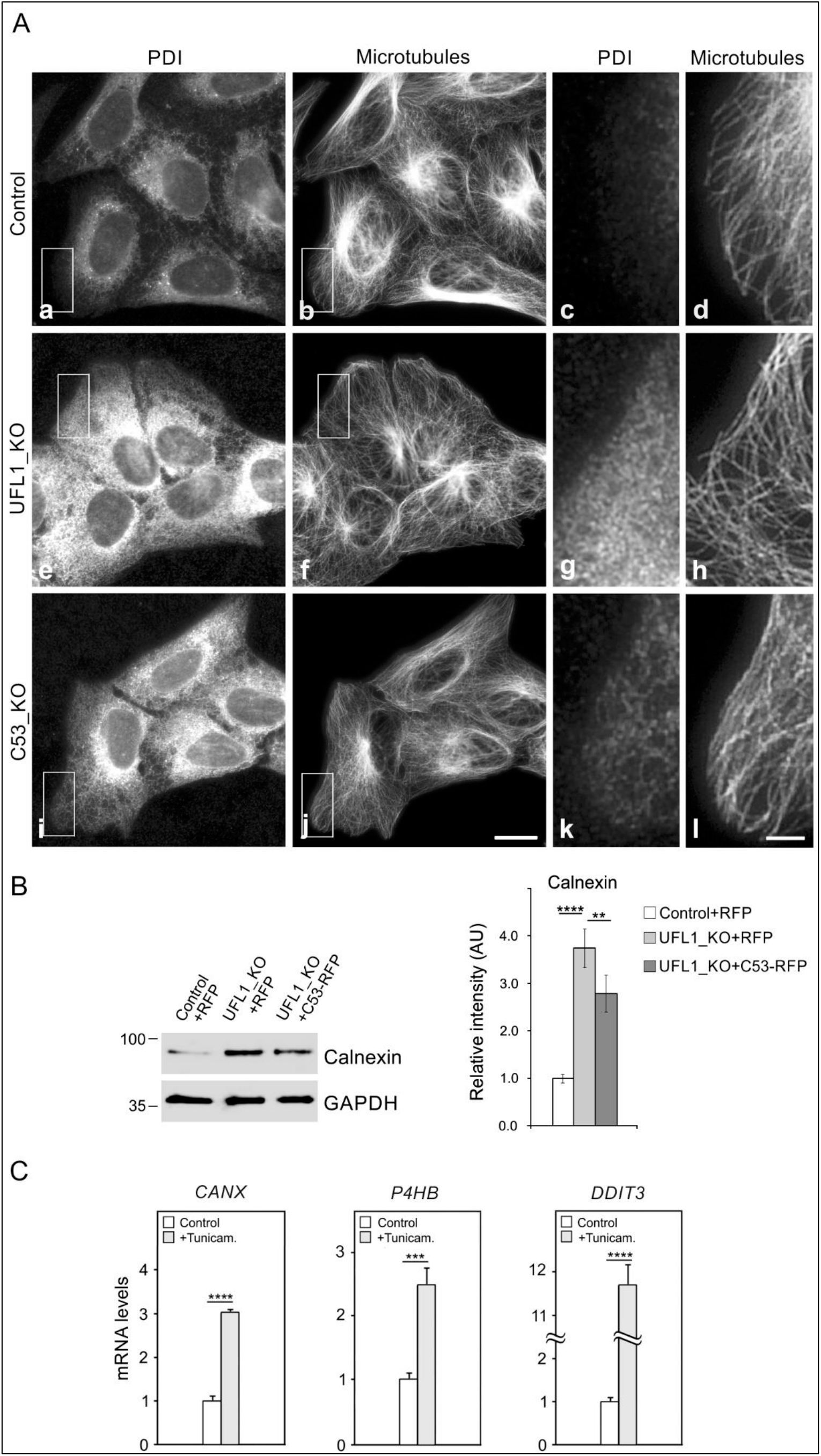
Deletion of UFL1 or C53 induces subcellular redistribution of PDI, and C53 attenuates calnexin expression in cells lacking UFL1. Tunicamycin causes transcriptional upregulation of ER stress-associated proteins. (**A**) Immunofluorescence microscopy. (**a-d**) Control, (**e-h**) UFL1-deficient (UFL1_KO) and (**i-l**) C53-deficient (C53_KO) U2OS cells. Cells were fixed and double-labeled for PDI (**a, e, i**) and β-tubulin (**b, f, j**; Microtubules). Higher magnification views of the regions delimited by rectangles are shown on the right of images from control (**c-d**), UFL1_KO (**g-h**), and C53_KO cells (**k-l**). Images (**a, e, i**) and (**c, g, k**) were collected and processed in exactly the same manner. Fixation F/Tx. Scale bars, 20 μm (**j**) and 5 μm (**l**). (**B**) Immunoblot analysis of a phenotypic rescue experiment. Whole-cell lysates from control cells expressing TagRFP (Control+RFP), UFL1_KO cells expressing TagRFP (UFL1_KO+RFP), and UFL1_KO cells rescued by C53-TagRFP (UFL1_KO+C53-RFP). Blots were probed with Abs to calnexin and GAPDH (loading control). Densitometric quantification of immunoblots is shown on the right. Relative intensities of corresponding proteins normalized to control cells and the amount of GAPDH in individual samples. Values indicate mean ± SD (n=4). One-way ANOVA with Sidak’s multiple comparisons test was performed to determine statistical significance. (**C**) Transcription of calnexin *(CANX),* PDI *(P4HB),* and DDIT3 *(DDIT3)* genes in cells treated with tunicamycin relative to the levels in untreated control cells. Data represent the mean ± SD (n=3). Two-tailed, unpaired Student’s *t* test was performed to determine statistical significance. **, p < 0.01; ***, p < 0.001; ****, p < 0.0001.

## List of source data

Figure 4 source data (Excel file). Underlying data for graphs in panels A, B and C.

Figure 5 source data (Excel file). Underlying data for graphs in panels A, B, D, E, G and H.

Figure 6 source data (Excel file). Underlying data for graphs in panels B, C, E, F, H and I.

Figure 7 source data (Excel file). Underlying data for graph in panel A.

Figure 8 source data (Excel file). Underlying data for graphs in panels B, C, D and F.

Figure 9 source data (Excel file). Underlying data for graphs in panels B, D and E.

Figure S3 source data (Excel file). Underlying data for graphs in panels A, B, C and D.

Figure S4 source data (Excel file). Underlying data for graphs in panels B and C.

